# Evidence for a core set of microbial lichen symbionts from a global survey of metagenomes

**DOI:** 10.1101/2023.02.02.524463

**Authors:** Gulnara Tagirdzhanova, Paul Saary, Ellen S. Cameron, Arkadiy I. Garber, David Díaz Escandón, Spencer Goyette, Veera Tuovinen Nogerius, Alfredo Passo, Helmut Mayrhofer, Håkon Holien, Tor Tønsberg, Lisa Y. Stein, Robert D. Finn, Toby Spribille

## Abstract

Lichens are the archetypal symbiosis and the one for which the term was coined. Although application of shotgun sequencing techniques has shown that many lichen symbioses can harbour more symbionts than the canonically recognized fungus and photobiont, no global census of lichen organismal composition has been undertaken. Here, we analyze the genome content of 437 lichen metagenomes from six continents, and show that four bacterial lineages occur in the majority of lichen symbioses, at a frequency on par with algal photobionts. A single bacterial genus, *Lichenihabitans*, occurs in nearly one-third of all lichens sampled. Genome annotations from the most common lichen bacterial symbionts suggest they are aerobic anoxygenic photoheterotrophs and produce essential vitamins, but do not fix nitrogen. We also detected secondary basidiomycete symbionts in about two-thirds of analyzed metagenomes. Our survey suggests a core set of four to seven microbial symbionts are involved in forming and maintaining lichen symbioses.

## Main

In some biological systems, unrelated organisms have evolved interactions so stable and integrated that they appear to function, to the casual observer, as one. This phenomenon was discovered over 150 years ago in lichens. Long thought to constitute a single organism, lichens were revealed to be a tightly integrated relationship of a fungus and phototrophic alga and/or cyanobacterium (collectively often called ‘photobionts’), using a combination of culturing experiments and microscopy^1^. The coexistence of these symbionts has long been explained as a function of the fungus using sugars produced by the carbon-fixing phototroph^1,2,3^, though the exact function of these sugars, and what the phototroph gets from the relationship, is debated^4,5^. The view of lichen symbiosis as a fundamentally two-partner system has changed little in 150 years, despite nearly 100 years of evidence from culturing, and later barcode sequencing, that lichen symbioses harbour rich communities of bacteria^6^ and additional fungi^7^. Such discoveries have prompted speculation that the identified microbes play a role in lichen metabolism^6,8^.

Shotgun RNA- and DNA-sequencing has revealed that some lichen symbioses consistently contain two or three fungal species^9,10,11^. However, these surveys have focused on only a few symbioses, and largely excluded prokaryotic diversity. Nonetheless, hundreds of metagenomes have been sequenced in recent years and are available in public databases. Most appear to have been generated for the purposes of retrieving the genome of the quantitatively dominant fungal symbiont, here referred to as lichen fungal symbiont (LFS). The potential of these metagenomes for interrogating organismal composition in lichens remains however largely unexplored. Here, we analyzed 412 publicly available and 25 newly generated lichen metagenomes. From the recovered metagenome-assembled genomes, we construct phylogenetic trees, map their occurrence across our dataset, and predict based on genome annotations how the most frequent microbial players integrate into the lichen metabolic system.

## Results

### Lichen metagenomes yielded 1000 genomes

We obtained a dataset of 437 lichen metagenomes, including nearly every published metagenome and 25 metagenomes generated *de novo* (Supplementary Table 1). This dataset included representatives of most major lichen groups (Extended Data Fig. 1). The majority (n=387) came from lichens in which the LFS was from the class Lecanoromycetes (Ascomycota). The rest came from lichens in which the LFS belongs to other ascomycete groups (Arthoniomycetes, Dothideomycetes, Eurotiomycetes, or Lichinomycetes). The metagenomes were generated from samples collected in various geographic locations on six continents (Extended Data Fig. 2).

We assembled each metagenome individually and binned them using CONCOCT^12^ and MetaBAT2^13^ (Fig. 1a). This resulted in 17,390 bins, which we screened to obtain metagenomic assembled genomes (hereafter genomes). To identify genomes and assess their quality, we used CheckM^14^ for prokaryotes and EukCC^15^ for eukaryotes. We retained only the bins that passed the quality threshold QS50^14^ (defined as completeness-5*contamination≥50). Following bin merging (see methods), this corresponded to 1142 genomes. Since the same genome can be obtained multiple times, we dereplicated the genome set with dRep^16^ to obtain species-level representatives, defined at 95% average nucleotide identity (hereafter ‘species’). The final set included genomes of 1000 species: 674 bacterial, 294 fungal, and 32 algal (Fig. 2, Supplementary Table 2). Of the 437 metagenomes, 375 had at least one recoverable genome; the derivation of this and all subsequently discussed subsets is explained in detail in the Supplementary Note.

**Fig. 1.**
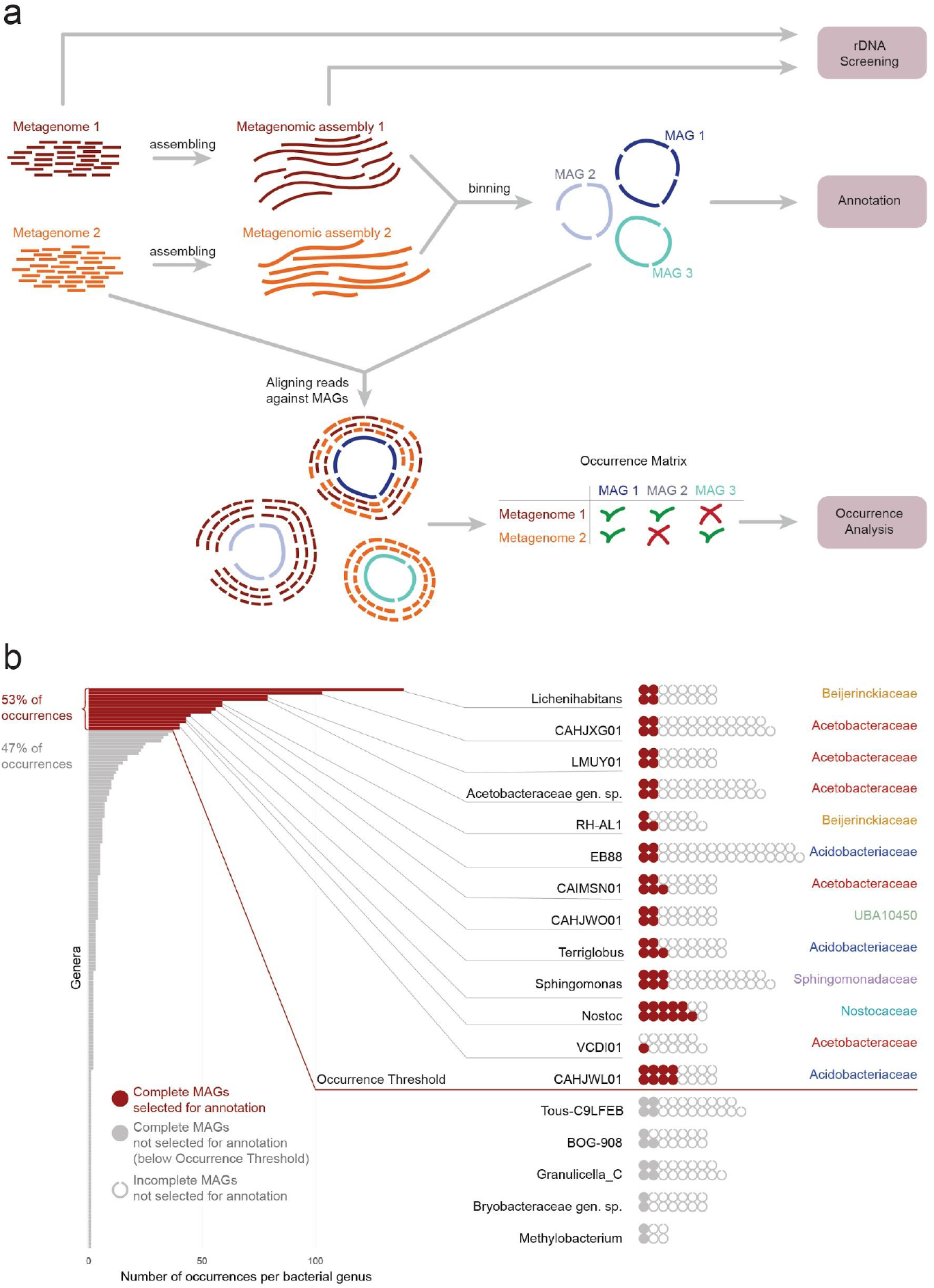
a. Flowchart of the bioinformatic analysis of this study. **b**. Selecting genomes for annotation. The bar graph on the left shows the total number of occurrences per bacterial genus, with genera listed in decreasing order. Red bars represent the genera selected for functional annotation, which were the 13 most frequent genera together accounting for 53% of all bacterial occurrences. The waffle graph on the right shows the most frequent bacterial genera and genomes assigned to them. Red circles represent genomes selected for annotation: all genomes from the selected genera that had >95% completeness. Full gray circles represent genomes with >95% completeness that belonged to less frequent genera and were not annotated. The incomplete gray circles represent genomes with ≤95% completeness, also not annotated.

**Fig. 2.**
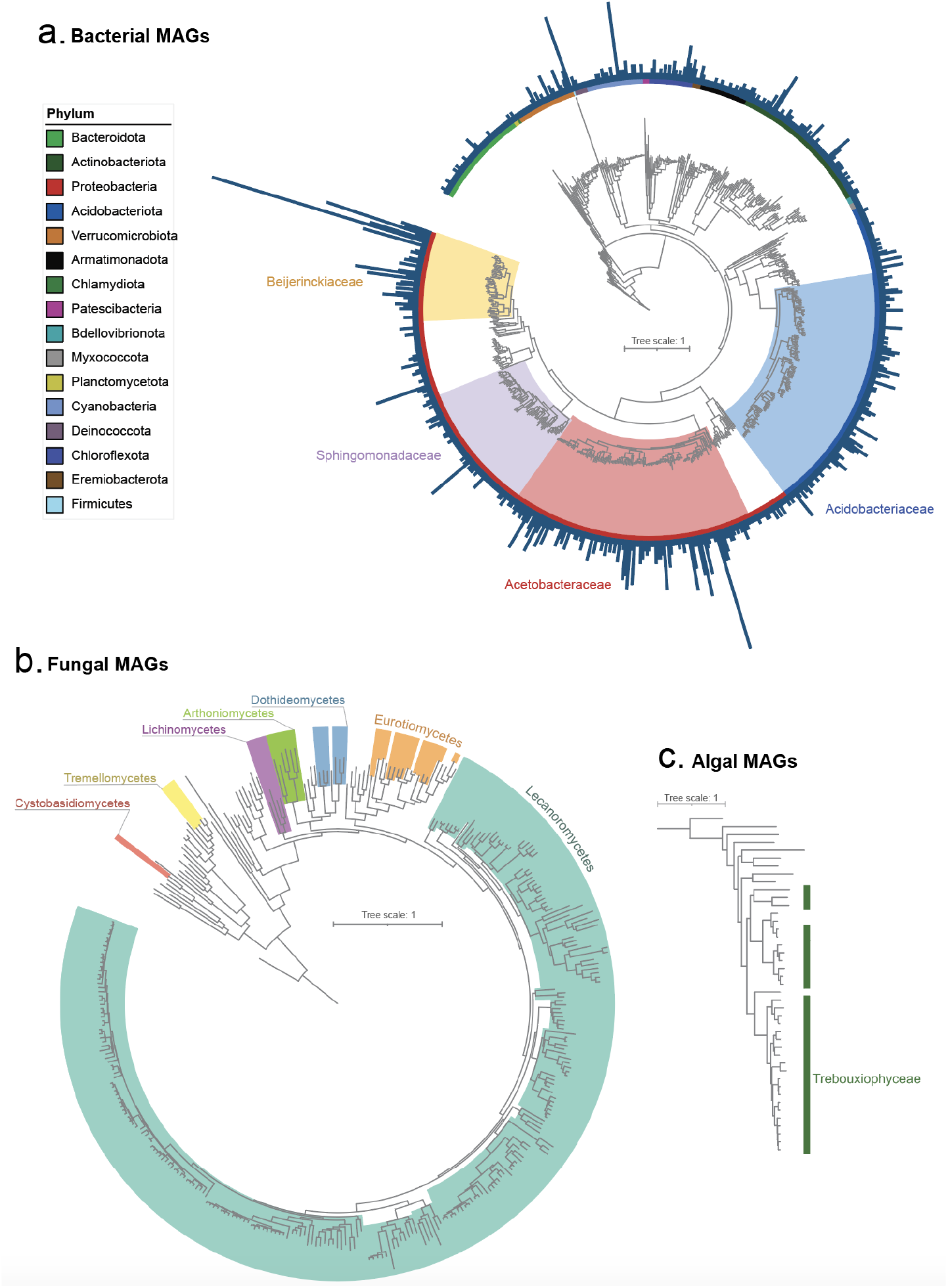
Maximum likelihood phylogenetic trees of the genomes. The trees are calculated using IQ-TREE and are based on alignments of marker genes (for prokaryotes: 120 marker genes from GTDB-Tk, for algae: 1296 genes from BUSCO, for fungi: 709 genes from BUSCO). The annotation tracks show the clades the genomes were assigned to, while the taxonomy of the reference genomes is not given. a. Tree of the prokaryotic genomes obtained from lichen metagenomes. The four most frequent bacterial families are highlighted. The bars represent the number of occurrences for each given genome. b. Fungal tree. c. Algal tree.

As expected, most eukaryotic genomes belonged to the two quantitatively dominant lichen symbionts, the LFS and the alga. All recovered algal genomes came from Trebouxiophyceae, Chlorophyta and belonged to known lineages of lichen symbionts. A total of 85% of fungal genomes came from known lineages of LFSs. The remaining fungi included two groups of basidiomycete fungi stably associated with lichens — Cystobasidiomycetes^9^ and Tremellomycetes^10^ — as well as non-LFS ascomycetes from the classes Dothideomycetes and Eurotiomycetes. Detected bacteria included representatives of 16 phyla (Fig. 2). Nearly 97% of detected bacterial species were unassignable to known species represented in GTDB, though most could be assigned to genera and families. The majority of bacterial species (51%) came from just four families: Acetobacteraceae, Beijerinckiaceae, Sphingomonadaceae (Alphaproteobacteria), and Acidobacteriaceae (Acidobacteriae) (Supplementary Table 2).

### Four bacterial families dominate lichen metagenomes

For each of the 1000 species identified above, we identified all instances in which that species was recovered as a genome passing threshold in the 375 metagenomes. This yielded a unique occurrence count per species (hereafter ‘occurrences’). Based on these counts, the same four bacterial families dominated, occurring in 64% of the metagenomes and together accounting for 47% of all occurrences and 62% of bacterial occurrences. These families even exceeded the occurrence counts of fungi (93% of metagenomes/20% of occurrences) and photobionts (algae: 17%/4%; Cyanobacteria: 13%/ 3%). One bacterial genus, *Lichenihabitans* (Beijerinckiaceae), accounted for 139 occurrences in 99 metagenomes (Supplementary Table 3), and a single *Lichenihabitans* species was detected in 52 metagenomes, including samples from different regions and in lichens involving distantly related LFSs.

We hypothesized that the recovery of complete genomes would correlate with sequencing depth, which varied widely. Indeed, greater sequencing depth translated into a higher number of genomes recovered from a metagenome (Extended Data Fig. 3. Supplementary Table 4). Moreover, in shallowly sequenced metagenomes, even the canonical LFS and algal genomes were below quality thresholds (Extended Data Fig. 4). Over 30% (n=89) of the metagenomes yielded no fungal genome that passed quality thresholds and 77% (n=335) yielded no photobiont genome, leaving a total of 330 metagenomes that yielded an LFS genome.

We reasoned that application of stringent thresholds for genome recovery would lead to undercounting organisms in the metagenomes. Accordingly, we screened for unambiguously assignable SSU rRNA gene sequences for all bacterial groups plus Trebouxiophyceae, Cystobasidiomycetes and Tremellomycetes, both in metagenomic assemblies and in sets of raw, unassembled reads. The most frequently detected organisms, in addition to the LFS and alga, were the four bacterial families identified in the metagenomic binning. Each of these four families was detected in over 60% of metagenomic assemblies and 90% of raw read sets (Fig. 3, Supplementary Table 5). The genus *Lichenihabitans* was found in 47% of metagenomic assemblies and 88% of read sets. Cystobasidiomycete and tremellomycete fungi were detected in 8% and 14% of assemblies and 30% and 62% of unassembled read sets, respectively.

**Fig. 3.**
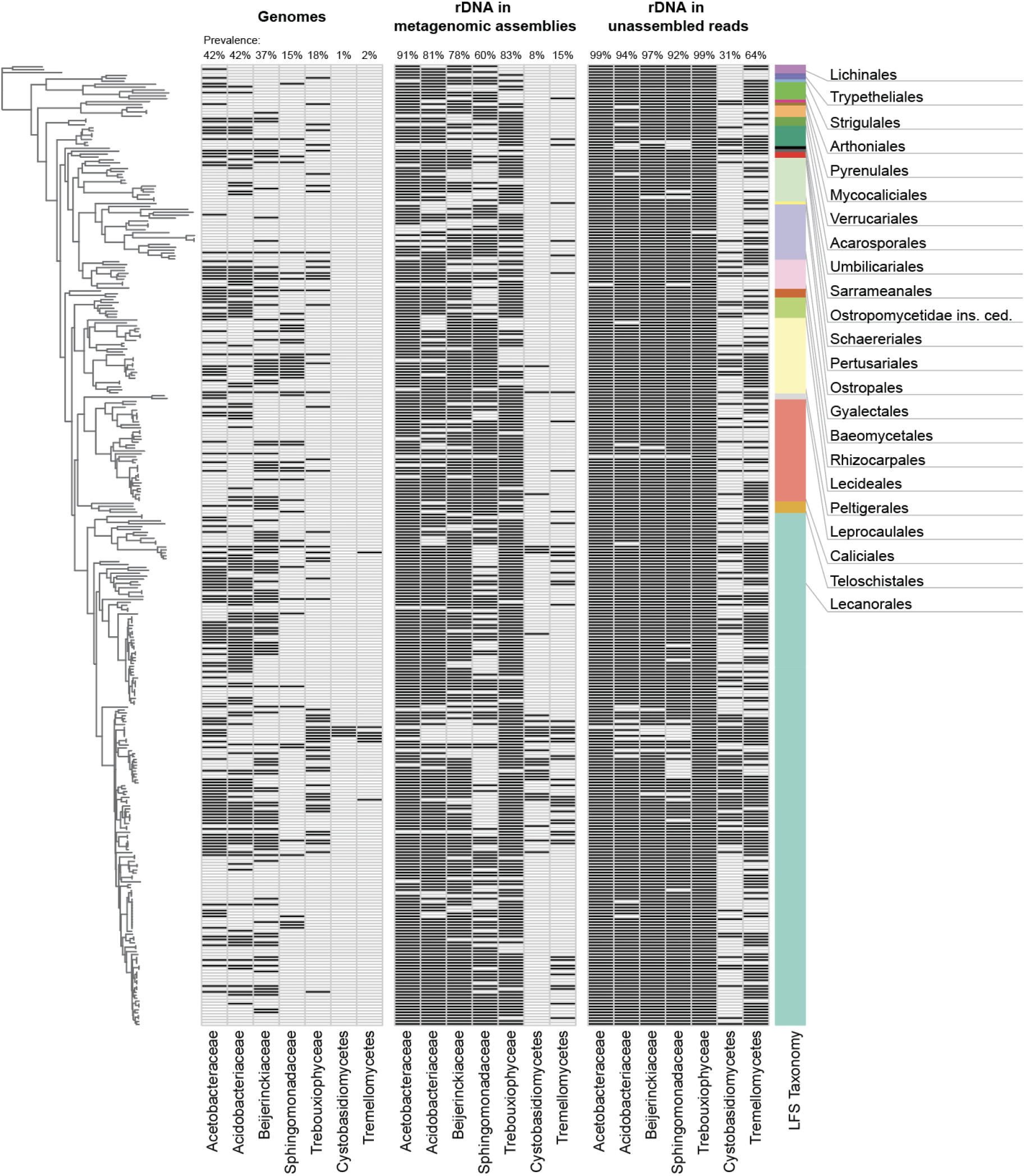
Detection of the key groups of symbionts in lichen metagenomes based on three methods of screening: presence of genomes, presence of SSU rRNA gene in the metagenomic assemblies, and presence of the SSU rRNA in the raw, unassembled reads. The tree on the left shows phylogeny of the LFS (each metagenome has only one LFS that also gives the name to the lichen). Here are shown data on the four most frequent bacterial families and on the three eukaryotic lineages known to be stably associated with lichens; co-occurrence with an LFS lineage is indicated by a dark stripe. Here we only show metagenomes that 1) yielded an LFS genome, and 2) are not labeled as deriving from potentially misidentified lichen samples (n=330; see Supplementary Data).

### Bacterial coverage often exceeds that of the alga

As a proxy for organismal cellular abundance, we calculated ratios of genome coverage depth (average number of reads per base) within each metagenome. Photobiont-to-LFS ratios ranged from 1:22 when the photobiont was an alga, to 2:1 when it was cyanobacterial. Even non-cyanobacterial bacteria achieved high coverage ratios. In 43% of occurrences, coverage depth of these bacterial lineages exceeded that of the alga. Similarly, individual bacterial lineages belonging to the four most frequent families often registered coverage depths comparable to those of the alga (median 1:1.4, ranging from 1:272 to 429:1). While coverage depth of individual bacterial lineages tended to be only a fraction of LFS coverage (median: 1:12, ranging from 1:402 to 6:1) (Extended Data Fig. 5, Supplementary Table 6), the combined coverage depth of non-cyanobacterial bacterial genomes exceeded LFS coverage depth in 30% of metagenomes.

### Patterns of bacterial occurrence

To assess whether detected organisms occurred randomly relative to the LFS, we mapped co-occurrence networks, beginning with the widely recognized algal symbionts. For these, the network formed a centroid with two “dominant” nodes, accounting for 31% of all algal occurrences, while the majority of lineages occurred only once (Fig. 4a). Bacterial networks differed depending on the family. In Acetobacteraceae and Beijerinckiaceae, many LFSs consorted with few lineages, forming network structures more centralized than the LFS-algal networks (Fig. 4b-d). Acidobacteriaceae, by contrast, exhibited decentralized networks similar to those of cyanobacteria, with few bacterial species associated with more than three LFSs.

**Fig. 4.**
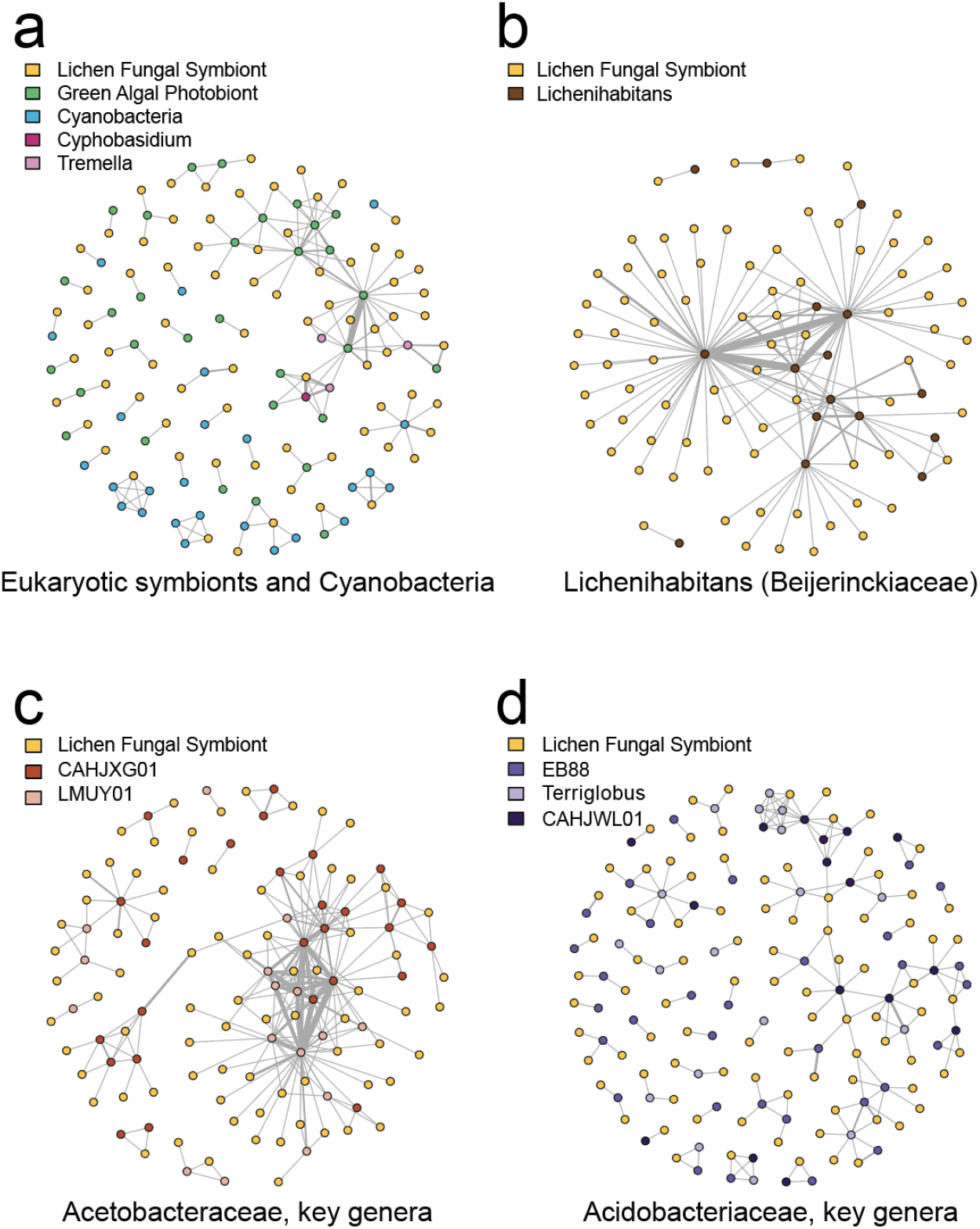
Co-occurrence networks of lichen symbionts based on presence-absence of genomes in each metagenome. Each node is a genome, and edges represent the co-occurrence of genomes within one metagenome; the thicker the edge, the more often two genomes co-occur. Nodes are coloured based on the taxonomy and function of the symbiont; in each network, yellow nodes represent genomes of the LFS. Only data on metagenomes that yielded an LFS genome are shown. **a**. Co-occurrence of LFSs, other known eukaryotic symbionts, and Cyanobacteria. **b**. Co-occurrence of LFSs and *Lichenihabitans*. **c**. Co-occurrence of LFSs and most frequent genera of Acetobacteraceae. **d**. Co-occurrence of LFSs and most frequent genera of Acidobacteriaceae.

While some bacterial species were present with high frequency across all samples, others were disproportionately associated with specific symbionts. For example, in lichens with an LFS from Lichinomycetes, the most frequent bacterial group was Chloroflexota (Supplementary Table 3). Similarly, lichens with cyanobacterial photobionts had more Sphingomonadaceae occurrences compared to those with only algal photobionts (Supplementary Table 3).

### The majority of lichen bacteria are heterotrophs

To select bacterial genomes for functional annotation, we ranked bacterial genera by the number of occurrences and selected the 13 most frequent genera, which together accounted for over half of all bacterial occurrences, 37% of bacterial lineages and 53% of bacterial occurrences. Next, we analyzed the genomes assigned to these genera and retained all that were ≥95% complete and ≤5% contaminated (Fig. 2b, Supplementary Table 7), for a total of 63 genomes.

Fourteen annotated genomes possessed a complete Calvin-Bensen-Bassham cycle, suggesting that they fix carbon (Fig. 5a). These included cyanobacterial photobionts, which are known autotrophs, and, unexpectedly, three species in Acetobacteraceae. All other bacterial genomes have annotations consistent with a heterotrophic lifestyle (Fig. 5a). None of these bacteria possessed alternative carbon fixation pathways^17^: reductive citrate cycle (KEGG module M00173), 3-hydroxypropionate bi-cycle (M00376), hydroxypropionate-hydroxybutyrate cycle (M00375), dicarboxylate-hydroxybutyrate cycle (M00374), Wood-Ljungdahl pathway (M00377), or the phosphate acetyltransferase-acetate kinase pathway (M00579).

**Fig. 5.**
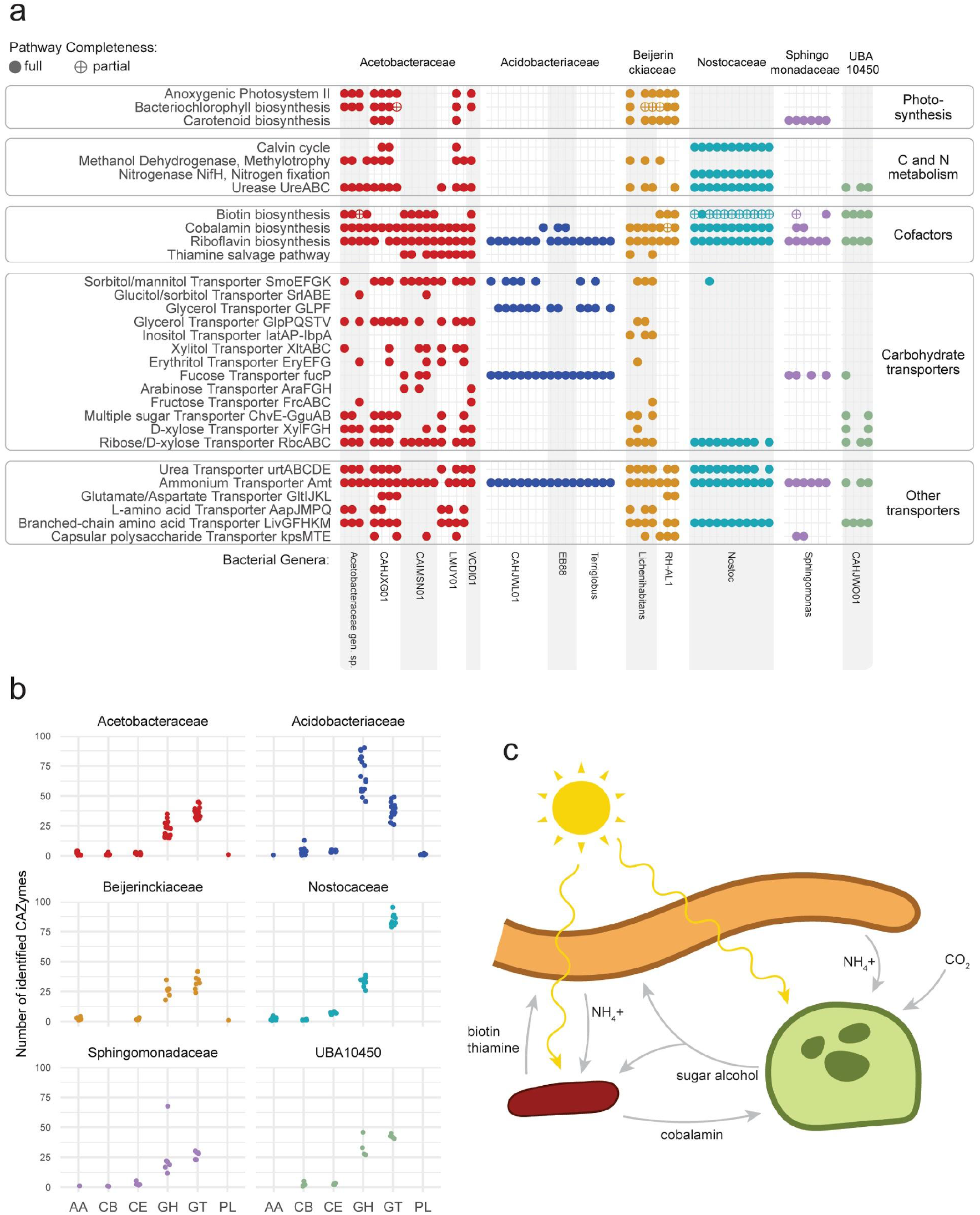
Functional annotation of bacterial genomes. **a**. Presence of selected pathways and protein complexes in the genomes of most common lichen bacteria. Each column represents one of the 63 annotated genomes, grouped by their taxonomy. We reconstructed pathways using KEGG; to detect carotenoid BGCs, we used antiSMASH. Here we show the presence of pathways and protein complexes potentially relevant to the symbiosis. For three pathways (biosynthesis of bacteriochlorophyll, biotin, and cobalamin), we also show partial completeness (allowing one missing gene). **b**. Number of genes assigned to each CAZy class per genome. We annotated CAZymes in the genomes selected for an in-depth annotation using dbcan. The data here are grouped on the family level. The CAZy classes are: Auxiliary Activities (AA), Carbohydrate-Binding Modules (CB), Carbohydrate Esterases (CE), Glycoside Hydrolases (GH), Glycosyl Transferases (GT), and Polysaccharide Lyases (PL). **c**. Reconstruction of the flow of metabolites in the lichen symbiosis, based on the functional annotations of genomes and evidence from the literature^85,86^.

To identify potential carbon sources for bacterial heterotrophs, we annotated transporter genes. Acetobacteraceae and one genus of Beijerinckiaceae (*Lichenihabitans*) have a large arsenal of transporters for various monosaccharides (ribose/D-xylose transporter *RbcABC*, D-xylose *XylFGH*, multiple sugars *ChvE-GguAB*, more rarely fructose *FrcABC* and L-arabinose *AraFGH*) and polyols (glycerol *GlpPQSTV*, sorbitol/mannitol *SmoEFGK*, xylitol *XltABC*, erythritol *EryEFG*, glucitol/sorbitol *SrlABE*) (Fig. 5a). Other studied bacterial families had fewer transporter systems, although Acidobacteriaceae had a glycerol transporter *GLPF*. Surprisingly, the one genus of Beijerinckiaceae (RH-AL1) did not possess any predicted sugar or polyol transporters.

Several lineages of Acetobacteraceae and Beijerinckiaceae exhibited annotations consistent with methylotrophy, the ability to use methanol as a carbon source (Fig. 5a). Methanol dehydrogenases (*XoxF* or *MxaF*) were detected in both Beijerinckiaceae genera with annotations (*Lichenihabitans* and RH-AL1), and five of six Acetobacteraceae genera (Fig. 5a), accounting for at least 75 occurrences in 50 metagenomes and as many as 499 occurrences in 187 metagenomes (if we assume that related genomes below completeness/contamination thresholds possess similar annotations). None of the studied bacteria appeared to use methane as none possess methane monooxygenase genes (Extended Data Fig. 6).

### The most frequent lichen bacteria are aerobic anoxygenic phototrophs

Almost a quarter of the annotated bacterial genomes (n=15), possessed both a complete set of anoxygenic photosystem II proteins (KEGG modules M00597 and M00165; *pufABCML-puhA*), and a bacteriochlorophyll biosynthesis pathway (*AcsF*, *ChlBNL*, *BchCFGPXYZ*). Many also contained carotenoid biosynthetic gene clusters (Fig. 5a). All of these genomes came from Acetobacteraceae and Beijerinckiaceae (including *Lichenihabitans*). This combination of pathways corresponds to a bacterial group known as aerobic anoxygenic phototrophs (AAPs^18^). The AAP profile is present in all Beijerinckiaceae genomes except one, and in half of Acetobacteraceae genomes. Conservatively, these genomes represent 84 occurrences in 56 metagenomes, and could represent as many as 499 occurrences in 187 metagenomes.

### Non-cyanobacterial lichen bacteria lack the *NifH* gene

Cyanobacteria in lichens fix nitrogen and supply it to other symbionts^19,20^, and it has been hypothesized that other lichen bacteria might do this as well^6^. Studies have reported PCR amplification of *NifH*, the key gene required for fixing nitrogen, from lichen samples that lack cyanobacteria^21,22^ and from bacteria isolated from lichens^23^. However, we did not recover *NifH* in any non-cyanobacterial genomes. To account for the possibility that *NifH* is present on a plasmid and therefore failed to be detected in genomes passing threshold, we searched for *NifH* across all metagenomic assemblies. Aside from cyanobacteria, we found only one Rhizobiales *NifH* in one metagenome.

By contrast, nearly all analyzed lineages possess the ammonium transporter *amtB* (Fig. 5a). In addition, in three families (Acetobacteraceae, Beijerinckiaceae and Nostocaceae), the majority of studied genomes had genes of the urea transporter *UrtABCDE* and urea metabolism *UreABC* (Fig. 5a). Most lineages possess various amino acids transporters: *LivGFHKM* for branched-chain amino acids, *AapJMPQ* for L-amino acids, and *GltIJKL* for glutamate/aspartate (Fig. 5a).

### Bacteria in lichens are predicted to produce cofactors essential for the eukaryotes

Lichen bacteria have been hypothesized to provide their eukaryotic partners with essential cofactors^6^. Several lineages possess pathways for biotin (vitamin B7) and thiamine (B1) biosynthesis, two vitamins required by the LFS (shown for several species^24,25^). Complete biotin biosynthesis pathways (KEGG modules M00123, M00577, and M00950) were present in 30% of analyzed genomes (n=18; Fig. 5a) accounting for at least 117 occurrences in 88 metagenomes and as many as 374 occurrences in 185 metagenomes. The thiamine biosynthesis pathway was present only partially, but the thiamine salvage pathway was complete in several Acetobacteraceae and Beijerinckiaceae (KEGG Module M00899 or *thiMDE*^26^).

Nearly all analyzed Acetobacteraceae, Beijerinckiaceae, and Nostocaceae genomes, present in at least 101 and as many as 236 metagenomes, encode pathways for biosynthesis of cobalamin (vitamin B12; KEGG Module M00122), required by many algae^27^. To establish whether lichen algae are cobalamin auxotrophs, we screened their genomes for the cobalamin-dependent methionine synthase gene *MetH*, as well as its cobalamin-independent alternative *MetE*. Most algal genomes (70%) possessed both *MetH* and *MetE* (Supplementary Table 8). This echoes results from Croft et al.^28^, who suggested that some algae preferentially use more efficient *MetH* in the presence of cobalamin, and switch to *MetE* only when cobalamin is absent.

### Lichen bacteria are not iron-limited

To test the hypothesis that lichen bacteria scavenge iron for the eukaryotic symbionts^29^, we profiled genes related to iron metabolism. Clusters potentially involved in siderophore biosynthesis were rare in all bacterial groups except Cyanobacteria. Even though every analyzed genome had genes related to siderophore transport (TonB-dependent receptors), these genes are not exclusively connected to, and cannot be viewed as evidence of, siderophore uptake^30^. Instead, the majority of genomes encoded iron ion transporters (Supplementary Table 9), which suggests that dissolved iron is present in the system and that symbionts are not iron-limited.

### Lichen bacteria can contribute to building and recycling lichen biomass

Among the 63 lineages selected for in-depth analysis, Acidobacteriaceae had twice as many glycoside hydrolases (GH) as the others (average=69 in Acidobacteriaceae, 22–34 in other families). Moreover, only in Acidobacteriaceae were GHs the dominant class of carbohydrate-active enzymes (CAZymes; Fig. 5b, Supplementary Table 10). Acidobacteriaceae possess several GH families targeting mannans: GH92 (mannosidase), GH125 (exo-alpha-1,6-mannosidase), GH38 (alpha-mannosidase), and GH76 (alpha-1,6-mannosidase/alpha-glucosidase). Also overrepresented in Acidobacteriaceae were GH18 (chitinase), GH55 (beta-1,3-glucanase), and GH51 (endoglucanase, endoxylanase, cellobiohydrolase) (Supplementary Table 11), all known to act on fungal polysaccharides.

In lichens, bacteria primarily inhabit the outermost layer, where they are embedded in an extracellular matrix^31^, thought to be synthesized by the LFS^32^. Our data suggest bacteria might contribute to it. Among annotated genomes, 29 encoded biosynthetic gene clusters with similarity to known exopolysaccharide-producing clusters (Supplementary Table 12; based on annotations with Jaccard distance <0.7 estimated by SanntiS, see Supplementary Note 1). Lineages with these clusters accounted for at least 102 occurrences in 61 metagenomes and as many as 697 occurrences in 228 metagenomes. In addition, genomes of several lineages from Acetobacteraceae, Beijerinckiaceae and Sphingomonadaceae encode a capsule polysaccharide transport system (Fig. 5a).

## Discussion

Our study represents a comprehensive, reference-free culture-independent census of lichen organismal components, taking inventory first of genomes that pass a quality filter, and then using these as queries to test for presence-absence across a range of sequencing depths. It draws from a broad swathe of lichen diversity across five ascomycete classes, all major lichen architecture types and six continents. Paradoxically, genome binning alone did not recover an alga or cyanobacterium as universally present in lichen symbioses, although we do not doubt that all material sequenced possessed at least one. This is easily explained by the wide range of sequencing depths in publicly available data, encompassing six orders of magnitude (from 70 Kbp to 35 Gbp per library), and the strong relationship between sequencing depth and the number of genomes passing QS50 thresholds (Extended Data Fig. 3). As a result, even well-known core symbionts such as algae only passed threshold in more deeply sequenced metagenomes. Indeed, even the LFS failed to pass threshold in 20% of metagenomes. A pitfall of sequencing depth is the false impression of organismal absence, as highlighted by our metabarcode screening.

Perhaps our most unexpected outcome was the global-scale frequency of specific bacteria taxa. All of the high-frequency bacterial families we report have been detected previously, either in amplicon^21,33,34^ or culturing studies^35^, but in only a few lichens. The most common bacterial genus, *Lichenihabitans* (syn. *Lichenibacterium*^35^, LAR-1^21^), had previously been reported in 25 lichen symbioses^21,33,34,35,36^. In the present study, a *Lichenihabitans* genome was recovered in 99 of 375 metagenomes and DNA detected in 88% of read sets. One family, Acetobacteraceae, was as frequent in our data (in 99% of raw read sets) as all representatives of trebouxiophycean algae taken together. Other families, including Beijerinckiaceae, Sphingomonadaceae and Acidobacteriaceae, far surpassed the detection frequency of basidiomycete yeasts of the classes Cystobasidiomycetes and Tremellomycetes, which were found in 33% and 60% of metagenomes, respectively. The network structuring of all bacterial groups except Acidobacteriaceae suggests extensive lineage sharing among lichen symbioses and mirrors the pattern exhibited by algae. Altogether, approximately 97% of species-level lineages we detected are novel to global databases.

Lichen bacteria have been postulated to play a key role in lichens, as integral ‘bacteriobionts’^37^ involved in nitrogen and carbon budgets. One of the oldest postulates, the fixation of atmospheric nitrogen^22,38^ by non-cyanobacterial bacteria, was not supported by our annotations, despite the frequency in our data of an order often associated with nitrogen fixation, Rhizobiales.

Whether the absence of *NifH* in our annotated Beijerinckiaceae genomes represents a loss during a shift to lichen environments remains to be tested. The suggestion that most non-cyanobacterial lichen bacteria are heterotrophs^37^ is supported by our annotations, except for some potential carbon fixers among Acetobacteraceae. Based on genome annotations alone, we cannot predict bacterial carbon sources, though many possess transporters consistent with use of the abundant polyols known to leak from lichens during wetting/drying cycles^39^. The capacity for methylotrophy, heretofore identified in bacterial annotations from a *Lobaria* lichen^40^, proved to be widespread. If demonstrated, use of methanol could further complicate calculations of the lichen carbon budget. An unexpectedly common feature in lichen bacteria only recently highlighted in one lineage by Pankratov et al.^41^ is their capacity for anoxygenic photosynthesis. It is impossible to state with certainty how many metagenomes contain AAP bacteria, as large differences in sequencing coverage in the 437 samples resulted in many genomes not passing threshold for inclusion in annotation. However, conservatively 17% and as many as 57% of metagenomes harboured AAPs. Originally discovered in aquatic environments^18,42,43^, AAPs have recently been identified in biological soil crusts^44,45^. Our data suggest that lichens, which like soil crusts undergo extreme desiccation and exposure, constitute a major global environment for AAPs. What functional role anoxygenic photosynthesis plays for heterotrophs in these environments is unknown.

A striking potential metabolic complementarity to emerge from our annotations is the capacity of many frequent lichen bacteria to code for cofactors needed by one of the dominant eukaryotic symbionts (Fig. 5c). Biomass accumulation in multiple LFSs have been shown to be dependent on biotin and thiamine^24,25^, cofactors predicted to be produced by a large percentage of the detected bacteria. The presence of genes and proteins associated with cofactor biosynthesis in the lichen *Lobaria*^40,46^ has been interpreted as evidence that bacteria support fungal metabolism and, in the case of cobalamin, sustain cobalamin-auxotrophic algae^6^. None of the algal genomes we queried are however unambiguously auxotrophic for cobalamin. Instead, they possess both *metE* and *metH*, suggesting that, if they function similarly to *Chlamydomonas reinhardtii*^28^, lichen algae might be facultatively dependent on cobalamin for methionine synthesis. Facultative dependence on cobalamin could be consistent with the habit of many lichen algae to spend a portion of their lifecycle outside of lichens^5^.

Much work remains before we gain a clear understanding of lichen metabolic flows. Our analyses suggest that a narrow cast of four to seven players, led by several high-frequency bacterial lineages, represent core microbial symbionts that service the eukaryotic partners while making a living in the lichen milieu. Lichen fungi and algae have always required vitamin supplements when grown *in vitro*; the metabolic toolboxes and ubiquity of lichen bacteria might provide a mechanism for covering these needs in nature. Resynthesis of whole lichens *in vitro* has proven difficult over the decades. Most successful resyntheses employ visually sorted lichen macerations, not axenic cultures in which bacterial cotransmission has been ruled out^47,48^. It remains to be seen if one or more of the bacterial strains identified here play a role in lichen synthesis. This may finally answer the question of what the minimum organismal components are that are required to make a lichen.

## Methods

Extended Methods and Results, Supplementary Figures and Tables are available as Supplementary Information.

### Dataset construction

We analyzed a total of 455 lichen metagenomes. Most of the data were obtained from NCBI (Supplementary Table 1), and 25 metagenomes were generated for this study (Supplementary Table 14). In the early stages of our analysis, we removed 43 metagenomes from NCBI, which were identified as duplicates (Supplementary Table 13, Supplementary Note). To generate new metagenomes, we collected lichen samples, froze them at –80°C and pulverized them using a TissueLyser II (Qiagen). We extracted DNA from the samples with DNAEasy Plant Mini Kit (Qiagen) and prepared metagenomic libraries. The libraries were sequenced on different Illumina HiSeq platforms to paired-end reads. The details on the procedure, including voucher information, library prep, and sequencing are given in Supplementary Table 14.

### Initial steps of metagenomic analysis

We started by assembling each metagenome individually and extracting metagenome-assembled genomes from them. The metagenomic libraries were filtered using fastp^49^ to remove adapters and low-quality bases, and the READ_QC module of the metaWRAP pipeline v.1.2^50^ to remove human contamination. The filtered data were assembled with metaSPAdes^51^. Individual assemblies were binned using CONCOCT^12^ and metaBAT2^13^. To refine prokaryotic genomes, we used the *binrefine* module of the metaWRAP pipeline and then evaluated all bins with CheckM v1.1.3^14^. Next, we selected all bins that passed the QS50 threshold and dereplicated them using dRep v3^16^ at 95% ANI (average nucleotide identity) and 30% AF (alignment fraction) thresholds in order to obtain species-level representatives.

We obtained taxonomic assignments for prokaryotic genomes using GTDB-Tk v1.5.0^52^, a tool based on the Genome Taxonomy DataBase (GTDB)^53^. In one case, we found an inconsistency between GTDB and the literature. *Lichenibacterium* and *Lichenihabitans* are two genera from Rhizobiales recently published within months of each other^35,36^. In GTDB, which provides taxonomic-rank normalization, *Lichenibacterium* is included in *Lichenihabitans*. We followed GTDB because it appears likely that the two studies independently described the same lineage. We generated a phylogenomic tree for all prokaryotic genomes that passed the QS50 threshold. For this tree, we used the marker gene alignment produced by GTDB-Tk (concatenated alignment of 120 loci). We generated the tree with IQ-TREE^54^ using the model finder (selected model: LG+F+R10) and 1000 bootstraps.

Eukaryotic genomes were identified and refined with EukCC v2^15^. Bins with a quality score of at least 50 were dereplicated with dRep at two levels: first, on the level of the individual binned metagenome (with the 99% ANI threshold); and second, on the level of the whole dataset, where bins from all metagenomes were dereplicated at 95% ANI and 40% AF to create species-representative genomes. For each genome, we calculated the EukCC and BUSCO5^55^ quality scores. To analyze the relationship between sequencing depth and recovery of genomes of the LFS and photobionts, we used the pre-dereplication genome set.

To obtain preliminary taxonomy annotations for eukaryotic genomes, we used BAT (CAT v5.2.3, database version: 20210107^56^), which predicts taxonomy based on searching predicted genes against the NCBI database. These taxonomic assignments were refined using phylogenomics. We separated all eukaryotic genomes into two groups: fungal and algal genomes. To both groups we added reference genomes (Supplementary Table 15). To compute the fungal phylogenomic tree, we used the Phylociraptor pipeline v0.9.6 (https://github.com/reslp/phylociraptor), executing the following steps (Supplementary Note). In each genome, we identified BUSCO universal single copy orthologs shared by at least 10% genomes in the set. We aligned the sequences and trimmed the alignments. From these data, we produced two trees: a coalescence tree that was reconstructed by ASTRAL v5.7.1^57^ from the individual gene trees (produced by IQ-TREE v2.0.7^54^), and a tree calculated from a concatenated alignment using IQ-TREE. We compared the coalescence and the concatenated phylogenies and reconciled detected discrepancies using signal base approximation in favor of the concatenated phylogeny. The algal phylogenomic tree was produced in the same way. The final trees were based on 1296 genes in algae and 709 genes in fungi.

### Occurrence analysis

To map genome occurrence across the metagenomes, we aligned reads from all metagenomes against all genomes using BWA-mem^58^. Next, we filtered the alignments using SAMtools^59^ to remove secondary alignments. All genomes that were at least 50% covered in a given metagenome were counted as present. Using these data, we constructed an occurrence matrix of genomes in metagenomes. To estimate the depth of coverage of genomes, we used the number of reads aligned to the genome, multiplied by the read length and divided by the total length of the contigs assigned to the genome.

In each metagenome, we identified the genome of the LFS. To do that, we manually inspected all fungal genomes present in a metagenome. If only one fungal genome was present, it was labeled as putative LFS. If multiple fungal genomes were present, we selected one as the LFS genome based on its position on the tree and the depth of coverage, since the genome of the main, most abundant LFS is expected to have greater depth of coverage than a genome from a fungal contaminant. To confirm the LFS assignments, we checked that the genome placement on the phylogenomic tree is consistent with the taxonomic assignment provided by the original data submitters in the NCBI metadata. If these did not match, we excluded metagenomes as potentially derived from misidentified samples. A total of 18 metagenomes were thus excluded (Supplementary Table 16). For all non-LFS genomes present in a metagenome, we estimated their relative abundances by computing their depth of coverage relative to that of the LFS genome. Only metagenomes that yielded an LFS genome were used for this analysis.

### rRNA gene-based screening

We searched metagenomic assemblies and raw, unassembled metagenomic data for the presence of the SSU rRNA gene of several lineages. This process consisted of two steps: the detection of 16S and 18S sequences, and their taxonomic assignment. 16S and 18S sequences were used for two reasons: first, they are the marker loci most frequently used for taxonomic profiling, and second, they tend to be present in multiple copies in a genome^60^ and therefore have better chances of being recovered in a shallow metagenome. For the first step, we used Metaxa2^61^, an HMM-based searching algorithm. For eukaryotic lineages, taxonomic placement was done through Metaxa2 as well. For bacteria, we used 16S sequences extracted by Metaxa2, to which we assigned taxonomic positions with IDTAXA^62^, which allowed us to use taxonomy consistent with GTDB.

### Functional analysis

We annotated all bacterial genomes using PROKKA v1.13^63^. Predicted proteins in lichen metagenomes were clustered to the MGnify protein database^64^ using the Linclust algorithm in mmseqs2 v13.45111^65^ at 90% coverage and 90% sequence identity. Next, we selected the genomes of the most frequent lineages and annotated them in depth. To select the genomes, we first ranked all bacterial genera based on the number of occurrences. For the genomes that did not have a genus level assignment, we used family-level annotations. Next, we selected the genomes assigned to the top 13 genera, and among them retained only genomes with a completeness score above 95% and contamination score below 10%, as estimated by CheckM.

For the selected genomes, we obtained functional annotations. We annotated predicted proteins against KEGG Orthology Database^66^ using KofamScan^67^. In addition, we used several tools to annotate groups of genes that are potentially relevant to the symbiotic lifestyle. We used following tools: run_dbcan (standalone tool of dbcan2, v3.0.2, https://github.com/linnabrown/run_dbcan) for annotations of Carbohydrate-Active EnZymes (CAZymes), FeGenie^30^ for the genes related to iron metabolism, and Sanntis (https://github.com/Finn-Lab/SanntiS) and antiSMASH v6.1.0^68^ for biosynthetic gene clusters. For *NifH*, we ran an additional tblastn search against the metagenomic assemblies and checked the taxonomy of the hits using reciprocal blast search against the NCBI database.

### Loss of function in Rhizobiales genomes

Rhizobiales genomes from our dataset lacked several functions typical for bacteria from this order. To put these genomes into the evolutionary context, we assembled a dataset that included 518 previously published genomes across the whole order^69^ (Supplementary Table 17) and a genome of *Rhodobacter* (GCF_009908265.2), which served as an outgroup. Using GTDB-Tk, we identified and aligned 120 marker genes. From this alignment, we generated a phylogenomic tree using IQ-TREE v2.1.2^54^. We used tblastn to screen all Rhizobiales genomes for the same genes related to nitrogen fixation, methanotrophy, and methylotrophy (Supplementary Table 18). For the genomes from GenBank, we confirmed that the tblastn results were consistent with the protein annotations available at NCBI.

### Data handling and visualization

Custom scripts used for data analysis and visualization were written in R v4.1.0^70^, using the following libraries: dplyr v1.0.8^71^, tidyr v1.2.0^72^, scales v1.1.1^73^, for data handling; ggplot2 v3.3.5^74^, ComplexHeatmap v2.11.1^75^, ape v5.0^76^, phangorn v2.8.1^77^, phytools v1.0-3^78^, circlize v0.4.14^79^, igraph v1.3.0^80^, qgraph v1.9.2^81^, treeio v1.16.2^82^, DECIPHER v2.14.0^83^, for data visualization. For visualizing phylogenetic trees, we also used iTOL^84^.

## Data availability

All genomes, annotations, and de novo generated raw data are submitted to ENA under the study accession PRJEB59037. Phylogenomic trees in Newick format are available at FigShare (doi.org/10.6084/m9.figshare.21913170.v1 and doi.org/10.6084/m9.figshare.21913212).

## Code availability

Custom scripts used for data analysis and visualization are available on GitHub (https://github.com/Spribille-lab/A-global-survey-of-lichen-symbionts-from-metagenomes) and FigShare (doi.org/10.6084/m9.figshare.21913221).

## Acknowledgments

GT acknowledges funding from Alberta Graduate Excellence Scholarship and Alberta Innovates Graduate Student Scholarship. Special thanks go to Piotr Łukasik for the help with producing *de novo* metagenomes and John McCutcheon for his support. This work was supported by an NSERC Discovery Grant to TS, and funding to TS through the Canada Research Chairs Program.

## Ethics information

The authors declare no competing interests.

