## Supplementary Material for "Evidence for a core set of microbial lichen symbionts from a global survey of metagenomes"

This documents contains:

- Supplementary Note: Extended Methods and Results
- Extended Data Figures
- Supplementary Figures
- Descriptions of Supplementary Tables

### Supplementary Note: Extended Methods and Results

The analysis was performed using published software and custom R scripts (R Core Team 2013). All generated code is available as a GitHub repository (https://github.com/Spribille-lab/Evidence-for-a-core-set-of-microbial-lichen-symbionts-from-a-global-survey-of-metagenomes/). The repository also contains: key results (figures and tables used in the manuscript and Supplementary Material), data used for generating tables and figures (phylogenetic trees and data tables), and a Readme file with a detailed description of how the analysis was performed. In addition, scripts used in the analysis and phylogenetic trees are available in FigShare repositories (doi.org/10.6084/m9.figshare.21913221, doi.org/10.6084/m9.figshare.21913170.v1, and doi.org/10.6084/m9.figshare.21913212).

### 1. Dataset construction

#### 1.1. Overview of the dataset

This study is based on 437 lichen metagenomes. The dataset included nearly all metagenomes available in NCBI and 25 additional metagenomes generated *de novo* (Supplementary Table 1). The dataset included lichens from across the diversity of lichen symbioses. It included lichens involving lichen fungal symbionts (LFS) from five different Ascomycota classes: Lecanoromycetes, Arthoniomycetes, Dothideomycetes, Eurotiomycetes, and Lichinomycetes. The dataset also covered the diversity of lichen photosynthetic symbionts (also known as photobionts). Lichen symbioses included in the analysis contained: a) chlorophyte algal photobionts, b) cyanobacterial photobionts, and c) both (Supplementary Table 1). Finally, our dataset included lichens with both main architecture types: macrolichens (three-dimensional architectures in the form of leafs and shrubs that are separated from the substratum) and crusts (biofilm-like architectures that adhere closely to the substratum or are even immersed in it).

Several metagenomes were initially selected for analysis and then removed on the ground of being duplicates of other metagenomes in the dataset. To identify such metagenomes, we used sourmash v4.2.2 (Pierce et al., 2019); we used the ‘sourmash sketch’ module to compute signatures and the ‘sourmash module’ to compare individual metagenomes. This way, we identified 43 pairs of identical metagenomes. All duplicates were shared between two studies, PRJNA731936 and PRJNA700635, suggesting that the same dataset was submitted to NCBI twice under different accession numbers. From each pair, we removed one metagenome (Supplementary Table 13). The set of 437 metagenomes used for the analysis (Supplementary Table 1) includes only unique metagenomes.

#### 1.2. *De novo* sequencing

To generate new metagenomes, we collected lichen samples, froze them at –80°C and pulverized them using a TissueLyser II (Qiagen). We extracted DNA from the samples with DNEasy Plant Mini Kit (Qiagen) and prepared metagenomic libraries. The libraries were sequenced on different Illumina HiSeq platforms to paired-end reads. The details on the procedure, including voucher information, library prep, and sequencing are given in Supplementary Table 14.

### 2. Metagenome assembly and binning

We started by assembling and binning each metagenome individually. The metagenomic libraries were filtered using fastp (Chen et al., 2018) to remove adapters and low-quality bases, and the READ_QC module of the metaWRAP pipeline v.1.2 (Uritskiy et al. 2018) to remove human contamination. The filtered data were assembled with metaSPAdes (Nurk et al. 2017). Individual assemblies were binned using CONCOCT (Alneberg et al. 2014) and metaBAT2 (Kang et al. 2019). The metagenomes that did not produce contigs sufficient for binning (n=20) were removed from the further analysis.

### 3. Taxonomic assignments of genomes

#### 3.1. Prokaryotic genomes

To obtain prokaryotic genomes, we refined all CONCOCT (Alneberg et al. 2014) and metaBAT2 (Kang et al. 2019) bins using the *binrefine* module of the metaWRAP pipeline (Uritskiy et al. 2018) and analyzed them using CheckM v1.1.3 (Parks et al. 2015). Bins that passed the QS50 threshold (n=738) were selected for further analysis. Next, we obtained species-level representatives by dereplicating the selected bins using dRep v3 (Olm et al. 2017) at 95% ANI (average nucleotide identity) and 30% AF (alignment fraction) thresholds. The final set of prokaryotic genomes included 674 species-level representatives (Supplementary Table 2).

We obtained taxonomic assignments for the species representative genomes using GTDB-Tk v1.5.0 (Chaumeil et al. 2020), a tool based on the Genome Taxonomy DataBase (GTDB). In one case, we found an inconsistency between GTDB and the literature, namely, *Lichenibacterium* and *Lichenihabitans.* These are two genera from Rhizobiales published within months of each other (Noh et al. 2019, Pankratov et al. 2020). In GTDB, which provides taxonomic-rank normalization, *Lichenibacterium* is included in *Lichenihabitans*. We followed GTDB because it appears likely that the two studies independently described the same lineage.

We computed a phylogenomic tree for the species representative genomes. For this tree, we used the marker gene alignment produced by GTDB-Tk (concatenated alignment of 120 loci). We generated the tree with IQ-TREE (Nguyen et al. 2015) using the model finder (selected model: LG+F+R10) and 1000 bootstraps.

#### 3.2. Eukaryotic genomes

Eukaryotic genomes were identified and refined with EukCC v2 (Saary et al. 2020). For that, we processed all bins from each run using the EukCC merging module, and screened the resulting bins to identify eukaryotic genomes. Bins with a quality score of at least 50 (n=407) were dereplicated with dRep (Olm et al. 2017) at two levels: first, on the level of the individual binned metagenome (with the 99% ANI threshold); and second on the level of the whole dataset, where bins from all metagenomes were dereplicated at 95% ANI and 40% AF to create species-representative genomes. The final set of eukaryotic genomes included 326 species-level representatives (Supplementary Table 2). For each genome, we calculated the EukCC and BUSCO5 (Seppey et al. 2019) quality scores.

To obtain preliminary taxonomy annotations for eukaryotic genomes, we used BAT (CAT v5.2.3, database version: 20210107; von Meijenfeldt et al. 2019), which predicts taxonomy based on the search against the NCBI database. According to BAT, all eukaryotic genomes belonged to either fungi, or chlorophyte algae. These BAT assignments were then refined using phylogenomics. We computed two phylogenomic trees, one for fungal genomes and the other for algae. To both genome sets we added reference genomes (Supplementary Table 15).

To compute the phylogenomic trees, we used the Phylociraptor pipeline (v0.9.9, https://github.com/reslp/phylociraptor). The pipeline included seven steps. First, we searched the genomes for 1519 universal single copy ortholog genes from the BUSCO set chlorophyta_odb10 (for algae) and 758 from fungi_odb10 (for fungi). We set a lower limit of 10% shared genes across all taxa: this way we could include even incomplete metagenome-assembled genomes. Second, we aligned the sequences using MAFFT (v7.464, Katoh & Standley 2013). Third, we trimmed the alignments using trimAl (v1.4.1, Capella-Gutiérrez et al. 2009); we kept only genes with more than 10 parsimony informative sites for a total of 1296 genes in algae and 709 genes in fungi. Fourth, we modeled the gene evolution and computed single-gene trees using IQ-TREE (v2.0.7; Nguyen et al. 2015). Fifth, we used the single-gene trees to reconstruct species trees with ASTRAL (v5.7.1; Zhang et al. 2018). Sixth, we concatenated all gene alignments into one, estimated the best substitution model for each gene, and computed a tree using IQ-TREE with 1000 ultra-fast bootstraps. Finally, we compared the ASTRAL-derived tree to the concatenated tree using the Phylociraptor utility for conflict estimation. No major discrepancies were recovered between the algal trees. The discrepancies between fungal phylogenies were reconciled using signal base approximation comparing site log-likelihoods calculated in IQ-TREE per topology, and using Phylogenetic signal parser (v1.1; Shen et al. 2017). Both genes and site signaling favored the concatenated tree over the coalescent. The trees in Newick formats are available at FigShare (doi.org/10.6084/m9.figshare.21913212).

The placement of eukaryotic genomes on the phylogenomic tree was consistent with our expectations, given what we knew about organisms participating in lichen symbioses. Algal genomes isolated from lichen metagenomes were all recovered in the class Trebouxiophyceae, split between Trebouxiales and the Elliptochloris clade. Fungal genomes included both ascomycetes (classes Arthoniomycetes, Dothideomycetes, Eurotiomycetes, Lecanoromycetes, and Lichinomycetes) and basidiomycetes (classes Cystobasidiomycetes and Tremellomycetes).

### 4. Occurrence analysis

#### 4.1. Constructing occurrence matrix

Each of the 1000 genomes recovered in sections 3.1 and 3.2 corresponds to a species-level lineage. To map the occurrences of the lineages across the analyzed samples, we followed the procedure from https://github.com/alexmsalmeida/metamap. First, we aligned reads from all metagenomes against all genomes using BWA-mem (Li & Durbin 2009) with default settings. Next, we filtered the mapped reads using SAMtools (Li et al. 2009) (samtools view -F 256 -uS). All genomes that were at least 50% covered in a given metagenome were counted as present.

Using these data, we constructed an occurrence matrix, where one lineage could occur in multiple metagenomes. In total, we recorded 2085 occurrences shared between 1000 lineages.

To estimate the depth of coverage of genomes in a given metagenome, we used the number of reads aligned to the genome, multiplied by the read length and divided by the total length of the contigs assigned to the genome.

**4.2. Assigning putative roles to genome occurrences**

Each lichen symbiosis contains the two quantitatively dominant partners: the LFS and the photobiont. The taxonomy of both these partners is known, with few exceptions in the case of the photobiont, for all described lichen symbioses. We used this knowledge to identify the genomes of the two dominant partners among the genomes present in a given metagenome. For the lichens that are known to have an algal photobiont, we identified the genomes assigned to Chlorophyta, and labeled them as photobiont. Similarly, for the lichens with cyanobacterial photobionts, we identified all cyanobacterial genomes.

To identify the LFS genomes, we inspected all fungal genomes present within a given metagenome. If only one fungal genome was present, it was labeled as putative LFS. If multiple fungal genomes were present, we labeled as putative LFS the fungal genome with the highest depth of coverage (since in lichens the LFS is expected to have greater cellular abundance than other fungi). The role was assigned to a genome occurrence, not a genome itself. In other words, we allowed fungal genomes to have different putative roles assigned to them in different metagenomes: for instance, one genome could be labeled as the LFS in one metagenome, and as a putative contaminant in others.

**4.3. Removing potentially misidentified samples**

For each metagenome, that yielded a putative LFS genome (n=348) we checked that its "content" (i.e. the position of the LFS genome on the phylogenomics tree) matches the "label" (i.e. the name of the lichen given in the NCBI metadata associated with the sample). In total, we identified 23 inconsistencies. In three metagenomes (SRR14722059, SRR14722135, SRR14722098), the inconsistency was easily resolved by correcting the putative LFS assignment, and giving it to a different fungal genome present in the metagenome. In additional two metagenomes (SRR14722289 and SRR14721950), NCBI metadata had inconsistencies within itself: the lichen name in the "organism" field did not match the name in the "library name". Our taxonomic placement agreed with the latter, and therefore we suspected that the "organism" field was filled in incorrectly during data uploading. We retained these two samples, correcting their names to their "library names".

The remaining 18 inconsistencies could not be resolved (Supplementary Table 16). We suspect that these metagenomes derived from misidentified specimens. For instance, *Rinodina brauniana* (SRR14722303) was expected to have an LFS from the order Caliciales. Instead, our phylogenomic analysis recovered its only putative LFS genome within the Lecanorales clade, together with the LFSs of *Lecanora* lichens. Given that *Rinodina* and *Lecanora* look somewhat similar, we suspected that the SRR14722303 metagenome was produced from a misidentified specimen. We excluded these metagenomes from some of the downstream analyses, where the identity of lichen symbiosis mattered (see below).

In total, 330 lichen metagenomes contained a confirmed LFS genome.

#### 4.5. Identifying most frequent bacterial groups

We ranked bacterial groups based on their frequency, defined as the total number of occurrences across the dataset. We summarized frequency on four taxonomic levels: species-level lineage, genus, family, and order. For the species-level lineages, we simply counted how many metagenomes they were detected in. For the higher taxonomic levels, we summed all occurrences of all lineages assigned to that group. If a genome did not have a genus level assignment, we used its family-level annotations (e.g. Acetobacteraceae gen. sp.). In addition, we ranked higher-level taxonomic groups based on how many species-level lineages from this group were detected. When calculating the percentage of metagenomes a given lineage was detected in, we only included metagenomes that yielded at least one genome (n=375).

On the level of bacterial families and orders both rankings largely agreed: the top four families were Acetobacteraceae, Beijerinckiaceae, Sphingomonadaceae, and Acidobacteriaceae. Together, these families accounted for 50% of all special-level bacterial lineages and 62% of all bacterial occurrences. On the level of genera, the 13 most frequent genera accounted for 53% of all bacterial occurrences. We visualized the number of occurrences per lineage (Fig. 2) using a custom script and iTOL (Letunic & Bork 2021).

To establish whether the composition of bacterial communities correlated with the LFS taxonomy, we compared the most frequent bacteria of different lichen groups. For that, we divided our dataset into 20 groups based on the taxonomy of the LFS. Lichens involving lecanoromycete fungi were split on the order level (Acarosporales, Baeomycetales, Caliciales, Gyalectales, Lecanorales, Lecideales, Leprocaulales, Peltigerales, Pertusariales, Rhizocarpales, Sarrameanales, Schaereriales, Teloschistales, Umbilicariales and the unassigned lineages of Ostropomycetidae *incertae sedis*); the remaining groups were defined on the class level (Arthoniomycetes, Dothideomycetes, Eurotiomycetes, Lichinomycetes). We excluded from this analysis metagenomes that did not yield a genome of the main fungal partner (n=27 metagenomes) or any bacterial genomes (n=80), and metagenomes produced from potentially misidentified samples (n=18).

For each lichen group, we created a list of the most common bacteria (Supplementary Table 3). The majority of the large groups shared the most common bacteria; this included lichens involving fungi from the orders Baeomycetales, Caliciales, Lecanorales, Pertusariales, and Teloschistales. Other groups exhibited a divergent bacterial composition. First, lichens involving LFSs from the Peltigerales (hereafter ‘peltigeralean fungi’) possessed Nostocaceae among the dominant bacteria, which was expected since cyanobacteria are the known primary or secondary photobionts in most lichens in this group (so-called “cyanolichens”). Less expected was the finding that Sphingomonadaceae were the second most common group in peltigeralean lichens, while being less frequent in other groups. Second, Burkholderiaceae were more common in both peltigeralean lichens and lichens with Umbilicariales as the main fungal symbiont than in other lichen groups. In addition, lichens with Umbilicariales fungi contained almost no Beijerinckiaceae genomes. Third, some lichen groups represented by a small number of metagenomes and/or shallowly sequenced metagenomes (e.g. involving fungi of the classes Arthoniomycetes or Lichinomycetes) exhibited high frequencies of bacteria otherwise rare in our dataset. However, this pattern might be an artifact caused by small sample sizes.

**4.6. Co-occurrence analysis**

To explore how lineages co-occur within lichen samples, we built co-occurrence network graphs (Fig. 4), using the occurrence matrix. We defined co-occurrence as an instance of two lineages occurring together in one metagenome. For this analysis, we focused on the groups that are known to stably occur in lichens (algae, Cyanobacteria, and Cystobasidiomycetes and

Tremellomycetes fungi), and on the most frequent bacterial groups (most frequent genera of Acetobacteraceae, Beijerinckiaceae, and Acidobacteriaceae). Only metagenomes that yielded an LFS genome were included in this analysis.

**4.7. rRNA gene-based screening**

In addition to profiling taxonomic composition of lichen metagenomes using genome presence/absence, we profiled SSU rRNA gene sequences detected in metagenomic assemblies and raw, unassembled metagenomic data. This process consisted of two steps: the detection of 16S and 18S sequences, and their taxonomic assignment. 16S and 18S sequences were used for two reasons: first, they are the marker loci most frequently used for taxonomic profiling, and second, they tend to be present in multiple copies in a genome (Gruber-Vodicka et al. 2020) and therefore have better chances of being recovered in a shallowly sequenced metagenome.

For the detection of rRNA gene sequences, we used Metaxa2 (Bengtsson-Palme et al. 2015), an HMM-based search algorithm. We created a custom Snakemake pipeline (Mölder et al. 2021) that used Metaxa2 to screen all metagenomic assemblies and unassembled read sets. The output produced by Metaxa2 included fasta files with the detected rRNA gene sequences, and their taxonomic assignments. We used these Metaxa2-derived classifications for the eukaryotic rRNA genes. For bacteria, we used the 16S sequences extracted by Metaxa2 and re-classified them using IDTAXA (Murali et al. 2018), which produced taxonomic assignments consistent with the classification used by GTDB.

We used the rRNA gene-based taxonomic profiles to screen metagenomes for the presence of seven groups of organisms: the four most frequent bacterial families (Acetobacteraceae, Beijerinckiaceae, Acidobacteriaceae, Sphingomonadaceae) and the three eukaryotic classes

(Trebouxiophyceae algae and Cystobasidiomycetes and Tremellomycetes fungi). For each group, we calculated its prevalence, defined as the number of metagenomes it was detected in. We calculated prevalence in two ways: one based on screening of metagenomic assemblies, the other based on screening of unassembled read sets. In all cases, the latter was higher than the former, and both were higher than the prevalence as suggested by the presence/absence of genomes. In this analysis, we only included metagenomes that yielded at least one genome (n=375), to remove metagenomes that were sequenced too shallowly to be representative.

Based on the rRNA gene screening, we computed the prevalence of each detected bacterial family (Supplementary Table 5). The four bacterial families that came out on top were the same families that had the highest number of genome occurrences: Acetobacteraceae, Beijerinckiaceae, Acidobacteriaceae, Sphingomonadaceae. These results were consistent between the screening of metagenomic assemblies and unassembled read sets.

**5. Analyzing relative abundance of symbionts**

We used the depth of coverage as a proxy for cellular abundance of organisms within lichen samples. To calculate relative abundance of a lineage in a sample, we calculated the depth of coverage ratio between the target genome and the LFS genome. Only metagenomes that yielded an LFS genome were included in this analysis. We visualized the relative abundances of several key groups: the 13 most frequent bacterial genera and the three eukaryotic groups known to be associated with lichens (algae, and Cystobasidiomycetes and Tremellomycetes fungi) (Extended Data Fig. 5).

We identified all samples in which the estimated total abundance of bacteria exceeds the abundance of the LFS. For that, we summed the depth of coverage per genome category in each metagenome individually. When calculating the total depth of coverage for bacteria, we excluded Cyanobacteria. This was done since their presence would skew the results: since in lichens Cyanobacteria often play a role as photobiont, their abundance tends to be much higher than that of other bacteria (Supplementary Table 6, Extended Data Fig. 5).

### 6. Sequencing depth vs Genome recovery

#### 6.1. Total number of genomes as a function of sequencing depth

To check how sequencing depth affects the taxonomic profiling of a metagenome, we plotted the number of genomes per metagenome as a function of sequencing depth. All metagenomes were used in this analysis, including those that did not yield any genomes.

The number of genomes per metagenome increased with increasing sequencing depth and did not appear to plateau (Extended Data Fig. 3). The highest number of genomes was 50 in the *Lobaria pulmonaria* metagenome (ERR4179390), which was also the second most deeply sequenced, with almost 35 Gbp of data. The number of genomes did not appear to depend on the architecture type of the lichen (crust vs. macrolichen).

High number of genomes in metagenomes was primarily driven by the bacterial genomes (Supplementary Table 4). Also, some metagenomes contained multiple fungal genomes, in addition to the LFS of the source lichen. Sometimes these extra fungal genomes belonged to LFSs of different, unrelated lichens. For example, a metagenome of *Platismatia glauca* (X3) contained two lecanoromycete genomes: One was most likely the LFS, as it was nearly identical to other *Platismatia* LFS genomes and had high coverage (221×). The other was low-coverage (4×) and nearly identical to the LFS from the *Hypogymnia physodes* (X14) metagenome.

#### 6.2. Recovery of the genomes of the main partners as a function of sequencing depth

We explored how sequencing depth affects the recovery of the two main partners, the LFS and the photobiont. We ran two versions of this analysis. In the first, we analyzed whether the symbiont genomes can be *detected* in a metagenome. In the second, we analyzed whether the symbiont genomes can be *isolated* from a metagenome *de novo*.

For the first version, we used the occurrence matrix created by aligning metagenomic reads against all isolated genomes (see section 4.1; only genomes that had the breadth of coverage >50% in a given metagenome were counted as present). We visualized the recovery of photobiont and LFS genomes as a function of sequencing depth (Extended Data Fig. 4). The minimal sequencing depth that allowed recovery of the LFS genome was about 250 Mbp, and at least 1.2 Gbp were required to obtain both the LFS and photobiont genomes (Extended Data Fig. 4). At the same time, several metagenomes with sequencing depths well above these thresholds failed to yield a genome for one or both main partners. This might be explained by higher complexity of those metagenomes or the presence of multiple strains, which would drive down the coverage depth for each genome.

For the second version, we analyzed the genomes produced during the metagenomic binning, including the genomes that were discarded during dereplication. We counted only genomes that were at least 90% complete according to EukCC. The goal of this analysis was to explore how much data is required to *de novo* produce high-quality genomes of the quantitatively dominant symbionts. As we expected, the bar for genome recovery was higher: The minimal sequencing depth that allowed recovery of the LFS genome was about 550 Mbp, and at least 2 Gbp were required to obtain both the LFS and photobiont genomes.

#### 6.3. Genome completeness as a function of coverage depth

We hypothesized that higher coverage depth results in a more complete genome assembly. To test that, we plotted the genome completeness (estimated by EukCC) against coverage depth. As expected, increasing coverage depth leads to increasing completeness (Supplementary Fig. 1). In general, a genome needs to have a >10x coverage depth to yield a high-quality (>90% complete) assembly during metagenomic binning.

Higher coverage depth also results in higher coverage breadth (Supplementary Fig. 1). To show this, we collected data on individual genome occurrences, i.e. for each case of a genome detected in a metagenome by aligning reads against the complete set of genomes. High coverage breadth (>90% of genome covered in reads) was achieved by genomes that had at least 5x coverage depth in this metagenome.

### 7. Functional annotations

#### 7.1. Protein space analysis

We annotated all bacterial genomes using PROKKA v1.13 (Seemann 2014). Predicted proteins were clustered to the MGnify protein database (Mitchell et al. 2020) using the linclust algorithm in mmseqs2 v13.45111 (Steinegger & Söding 2017) at 90% coverage and 90% sequence identity.

#### 7.2. Functional clustering

We explored the link between functional profiles of bacteria and their taxonomy. In bacteria, taxonomy and function do not correlate perfectly, and therefore some patterns can be lost in a taxonomy-based profiling. Could it be that multiple taxonomic groups, each too rare to attract our attention on its own, together form a single functional group that plays an important role in lichens? To test for this possibility, we a) predicted metabolic traits based on the genome annotations, and b) clustered genomes based on the presence/absence of these traits.

First, we obtained functional annotations for all bacterial genomes and used them for predicting metabolic traits. For that, predicted protein models were annotated against KEGG Orthologue Database (Kanehisa et al. 2002) using KofamScan (Aramaki et al. 2020). From the KEGG annotations, we inferred the presence/absence of KEGG modules, which represent distinct functional capabilities (e.g. a biosynthetic pathway, a transport system, etc.). To produce module annotations, we selected high-quality genomes (CheckM completeness score > 90%) and used their KEGG annotations to reconstruct KEGG modules with a script from Zoccarato et al. (2022). To compensate for potential false absences (e.g. caused by genes being split between contigs due to imperfect metagenome assembly), we allowed one missing gene per module.

Next, we clustered genomes based on the presence/absence of KEGG modules. For that, we constructed correlation matrices for metagenomes and for bacterial genera based on Pearson coefficients. We used the R library simplifyEnrichment (Gu & Hübschmann 2021) to compare different clustering methods: hdbscan, apcluster, MCL, walktrap, kmeans, binary_cut, and dynamicTreeCut. To estimate the taxonomic coherence of different clustering outcomes, we followed Zoccarato et al. (2022). The resulting clustering was visualized using the ComplexHeatmaps library (v2.11.1, Gu et al. 2016).

The three clustering methods that produced clustering with a reasonable number of clusters were: apcluster, hdbscan, and kmeans (Supplementary Fig. 2). The clusters produced by hdbscan closely followed the taxonomy, and each cluster, with two exceptions, mapped to one bacterial phylum (Supplementary Fig. 3). Apcluster and kmeans disagreed with taxonomy more often

(Supplementary Fig. 3). However, in the cases where the clustering was not consistent with taxonomy, apcluster and kmeans also did not agree with each other. For example, in both methods Proteobacteria were split between six clusters, but the boundaries of the clusters differed. We concluded that lichen bacteria cannot be separated into functional clusters beyond their taxonomic profiles.

**7.3. Selecting bacterial genomes for in depth annotation**

From bacterial genomes we selected the genomes of the most frequent lineages and annotated them in depth. To select the genomes, we first ranked all bacterial genera based on the number of occurrences. For the genomes that did not have a genus level assignment, we used family-level annotations. Next, we selected the genomes assigned to the top 13 genera, which together accounted for 53% of all bacterial occurrences. Finally, we filtered the genomes and retained only genomes with a completeness score above 95% and contamination score below 10%, as estimated by CheckM. The resulting set included 63 genomes from 13 genera and 6 families: Acetobacteraceae, Beijerinckiaceae, Acidobacteriaceae, Sphingomonadaceae, Nostocaceae (Nostocales, Cyanobacteria), and UBA10450 (Chthoniobacterales, Verrucomicrobia). (Supplementary Table 7).

**7.4. In-depth functional annotations**

We obtained functional annotations for all bacterial genomes and used them for predicting metabolic traits. We used the KEGG annotations (see 7.2), which were visualized using the KAAS webserver (Moriya et al. 2007). We focused on several metabolic traits, which we expected to be relevant to the symbiosis:

- Carbon metabolism. We screened the genomes for known carbon fixation pathways, including Calvin-Bensen path (KEGG module M00165), and six alternative pathways (see Assié et al. 2020): reductive citrate cycle (M00173), 3-hydroxypropionate bi-cycle (M00376), hydroxypropionate-hydroxybutylate cycle (M00375), dicarboxylate-hydroxybutyrate cycle (M00374), Wood-Ljungdahl pathway (M00377), or the phosphate acetyltransferase-acetate kinase pathway (M00579). We also searched for the genes related to C1 metabolism: methanol dehydrogenase (KEGG family K23995) and methane monooxygenase (K10946 and K16157).
- Nitrogen metabolism. We screened the genomes for the presence of nitrogenase *NifH* (K02588; involved in nitrogen fixation), and urease (EC 3.5.1.5).
- Photosynthesis. We screened the genomes for the proteins of Anoxygenic Photosystem II (M00597 and M00165; *pufABCML-puhA*), and for biosynthetic pathways for the photosynthetic pigments: bacteriochlorophyll (K04035, K04037, K04038, K04039, K11333, K11334, K11335, K11336, K11337, K04040, K10960; *AcsF*, *ChlBNL*, *BchCFGPXYZ*), and carotenoids (K02291, K10027, K09844, K09844, K09845, K09846).
- Transport systems. We annotated the following transport systems and transporters: sorbitol/mannitol transporter (K10227, K10228, K10229, K10111; *SmoEFGK*), urea transporter (K11959, K11960, K11961, K11962, K11963; *urtABCDE*), erythritol transporter (K17202, K17203, K17204; *EryEFG*), xylitol transporter (K17205, K17206, K17207; *XltABC*), inositol transporter (K17208, K17209, K17210; *IatAP-IbpA*), glycerol transporter (K17321 K17322, K17323, K17324, 17325; *GlpPQSTV*), fucose transporter (K02429; *FucP*), glycerol aquaporin transporter (K02440; *GLPF*),

glucitol/sorbitol transporter (K02781, K02782, K02783; *SrlABE*), ammonium transporter (K03320; *Amt*), ribose transporter (K10439, K10440, K10441; *RbcABC*), xylose transporter (K10543, K10544, K10545; *XylFGH*), multiple sugar transporter (K10546, K10547, K10548; *ChvE-GguAB*), fructose transporter (K10552, K10553, K10554; *FrcABC*), arabinose transporter (K10537, K10538, K10539; *AraFGH*), branched-chain amino acid transporter (K01999, K01997, K01998, K01995, K01996; *LivGFHKM*), L-amino acid transporter (K09969, K09970, K09971, K09972; *AapJMPQ*), glutamate transporter (K10001, K10002, K10003, K10004; *GltIJKL*), capsular transporter (K10107, K09688, K09689; *KpsMTE*).

- Cofactors. We searched for the biosynthetic pathways of the following cofactors: biotin (M00123, M00577, and M00950), thiamine (M00899; thiamine salvage pathway), cobalamin (M00122), and riboflavin (M00125).

To account for the possibility that *NifH* is present on a plasmid and therefore failed to be included in the genome during binning, we further searched *NifH* across all contigs of all metagenomic assemblies, using tblastn (Altschul et al. 1990) and a *NifH* sequence from NCBI as a query (Supplementary Table 18). We extracted all hits and checked their taxonomy using reciprocal blast search against the NCBI database using getLCA (https://github.com/frederikseersholm/getLCA). All hits were assigned to Cyanobacteria, with one exception: one metagenome (SRR14722280) contained a Rhizobiales hit on a low-coverage contig not assigned to any genome isolated from this metagenome.

To annotate the Carbohydrate-Active EnZymes (CAZymes), we used run_dbcan — a standalone tool of dbcan2 (v3.0.2, https://github.com/linnabrown/run_dbcan). Run_dbcan assigns predicted proteins to CAZy families, using two search methods: DIAMOND and HMMER. To create consensus annotations from the two sets of predictions, we followed Krüger et al. (2019): we retained only hits with e-value < 1e-20, >30% identity for the DIAMOND hits, and >30% query coverage for the HMMER hits; and only retained annotations that were consistent between the two methods.

We used FeGenie (Garber et al. 2020) to annotate the genes related to iron metabolism.

We used antiSMASH (v6.1.0, Blin et al. 2021) and SanntiS (v0.2.3, https://github.com/Finn-Lab/SanntiS) for annotating biosynthetic gene clusters (BGCs). Both tools predict BGCs from the genome sequences and annotate predicted BGCs by comparing them to the MiBIG database. We screened these annotations for two groups of interest: BGCs potentially producing carotenoids and BGCs potentially producing extracellular polysaccharides. While analyzing the antiSMASH results, we used only BGCs that had significant hits to BGCs from the MiBIG database according to the outputs of the KnownClusterBlast module. While analyzing the SanntiS results, we retained only annotations that had Jaccard distance score below 0.7. Our SanntiS results should be considered as preliminary, since they are based entirely on sequence similarity, and we did not further validate the annotations by taking into account the position of the domains or their copy number.

### 8. Loss of function in Rhizobiales

#### 8.1. Rhizobiales phylogenomic analysis

Rhizobiales genomes from our dataset lacked several functions typical for bacteria from this order. To put these genomes in evolutionary context, we computed a phylogenomic tree based on all Rhizobiales genomes from our dataset (n=58) and 518 reference genomes (taxon sampling following Volpiano et al. 2021; Supplementary Table 17), plus a genome of *Rhodobacter* (GCF_009908265.2), which served as an outgroup. Using GTDB-Tk (v1.5.0, Chaumeil et al. 2020), we identified and aligned 120 marker genes. From this alignment, we generated a phylogenomic tree using IQ-TREE (v2.1.2, Nguyen et al. 2015). The tree in the graphic and Newick formats is available at FigShare (doi.org/10.6084/m9.figshare.21913170.v1).

#### 8.2. Screening for genes related to nitrogen fixation and C1 metabolism

First, we screened the genomes for nitrogenase *NifH*. For that, we searched genomic assemblies (i.e. nucleotide FASTA files) using tblastn (Altschul et al. 1990) as described above. To verify that this approach was robust, we ran two additional searches only on NCBI-derived genomes: 1) we used blastp to search protein FASTA files, and 2) we screened the functional annotations provided by NCBI for the genes annotated as nitrogenases. All three searches (tblastn against nucleotide assemblies, blastp against protein predictions, and text search of NCBI annotations) gave consistent results.

Next, we used tblastn to screen the genomes for four genes associated with C1 metabolism: methane monooxygenases *PmoC* and *MmoX*, and methanol dehydrogenases *XxoF* and *MxaF* (Supplementary Table 18). The identity of the resulting hits was confirmed by a reciprocal search against the NCBI database. We combined all search results, and visualized the phylogenomic tree using iTOL (Letunic & Bork 2021).

### 9. Searching for cobalamin-dependent genes in algal genomes

To establish whether the algae in our dataset are cobalamin auxotrophs, we followed Croft et al. (2005), and screened the algal genomes for a cobalamin-dependent methionine synthase *MetH* and another gene performing the same function — cobalamin-independent *MetE*. First, we selected algal genomes with completeness >90% according to EukCC (n=19). Next, we screened them using tblastn (Altschul et al. 1990) and two protein sequences as a query (BAU71143.1 for *MetH* and BAU71146.1 for *MetE*). We confirmed the identity of the resulting hits by a reciprocal search against the NCBI database.

### Description of Supplementary Tables

**Supplementary Table 1.** Metagenomic data used in the analysis.

**Supplementary Table 2.** Metagenome-assembled genomes generated from the lichen metagenomic data. For prokaryotic genomes, contamination and completeness scores are based on the CheckM results, and taxonomic assignments are based on GTDB. For eukaryotes, contamination and completeness scores are based on the EukCC results, and taxonomic assignments are based on the BAT assignments corrected via phylogenomic analysis.

**Supplementary Table 3.** Most frequent bacterial lineages (on the family and genus level) in various groups of lichen metagenomes. Bacterial lineages were ranked based on the total number of their occurrences. For each lichen group here we give top-ten bacterial taxa.

**Supplementary Table 4.** Number of genomes from different categories in each metagenome. The genome categories were based on a combination of taxonomy and function. For the metagenomes that were identified as potentially derived from misidentified samples, we assigned their LFS to a separate category "Main Fungal Partner misassigned".

**Supplementary Table 5.** Bacterial families ranked on their prevalence in lichen metagenomes, as detected via rDNA-based screening in metagenomic assemblies and unassembled reads. Prevalence is calculated using only metagenomes that yielded at least one genome (n=75).

**Supplementary Table 6.** Median relative coverage depth of bacterial genomes. For each bacterial genome in each metagenome, we calculated relative coverage by dividing their coverage depth by the coverage depth of the LFS. Metagenomes that did not yield a genome of the LFS were excluded from this analysis.

**Supplementary Table 7.** Genomes selected for the in depth functional annotation. To select these genomes, we first calculated the number of occurrences per genus, and selected the 13 top genera. Next, we selected among them genomes with >95% completeness and <5% contamination, according to CheckM.

**Supplementary Table 8.** Presence/absence of cobalamin-dependent *MetH* and cobalamin-independent *MetE* in the high-quality algal genomes (completeness > 90%, contamination < 10%, according to EukCC).

**Supplementary Table 9.** Number of iron-related genes annotated with FeGenie.

**Supplementary Table 10.** Median number of CAZymes assigned to different classes per genome for the most frequent bacterial genera.

**Supplementary Table 11.** Number of CAZymes assigned to different CAZy families in the 63 genomes selected for in-depth annotation.

**Supplementary Table 12.** Biosynthetic Gene Clusters (BGCs) predicted for the selected 63 bacterial genomes. We used SanntiS to predict the BGCs and to annotate them by identifying the most similar BGC included in the MiBIG database. Only the hits with <0.7 Jaccard distance are listed in this table. BGCs that are similar to known clusters producing exopolysaccharides are indicated in bold.

**Supplementary Table 13.** Pairs of identical metagenomes (sourmash similarity=1) identified during analysis. Using sourmash, we identified pairs of identical metagenomes, potentially deriving from one dataset having been submitted to NCBI twice under different accession numbers. This table contains all such pairs, and shows which of the pair was retained for the further analysis.

**Supplementary Table 14.** Details on the metagenomes generated *de novo* for this study.

**Supplementary Table 15.** List of reference genomes used for construction of the phylogenomic trees.

**Supplementary Table 16.** Metagenomes derived from potentially misidentified samples. Metagenomes in this list had inconsistencies between their metadata and the taxonomy of the

LFS as estimated from the phylogenomic tree. These metagenomes were excluded from the occurrence analysis.

**Supplementary Table 17.** Reference genomes from NCBI used for the Rhizobiales phylogenomic tree. Taxon sampling followed Volpiano et al. 2021. Taxonomic classification was done using GTDB-Tk.

**Supplementary Table 18.** Sequences used as a blast query for the screening genomes for the signs of nitrogen fixation, methanotrophy, and methylotrophy.

### Extended data figures

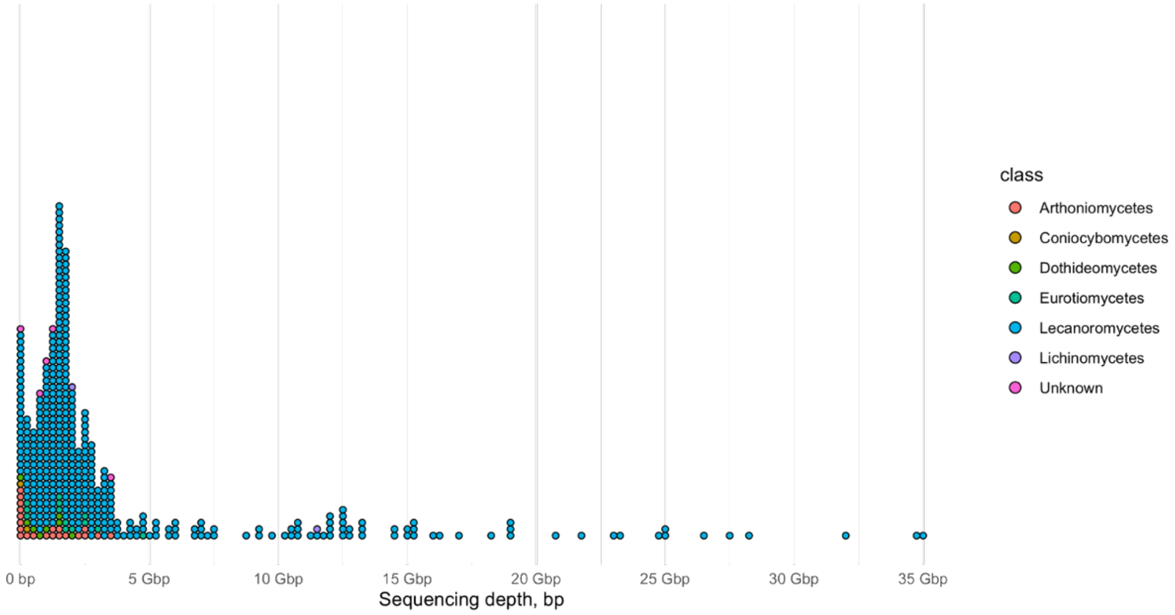


**Extended Data Fig. 1. Dot histogram of lichen metagenomes arranged by sequencing depth.** Each metagenome is shown with a dot; the dots are coloured according to the taxonomy of the LFS. The dots are arranged on the x-axis based on the sequencing depth (bp), and are "stacked" on top of each other. The peak in 0 bp represents all metagenomes with sequencing depth less than 250 Mbp.

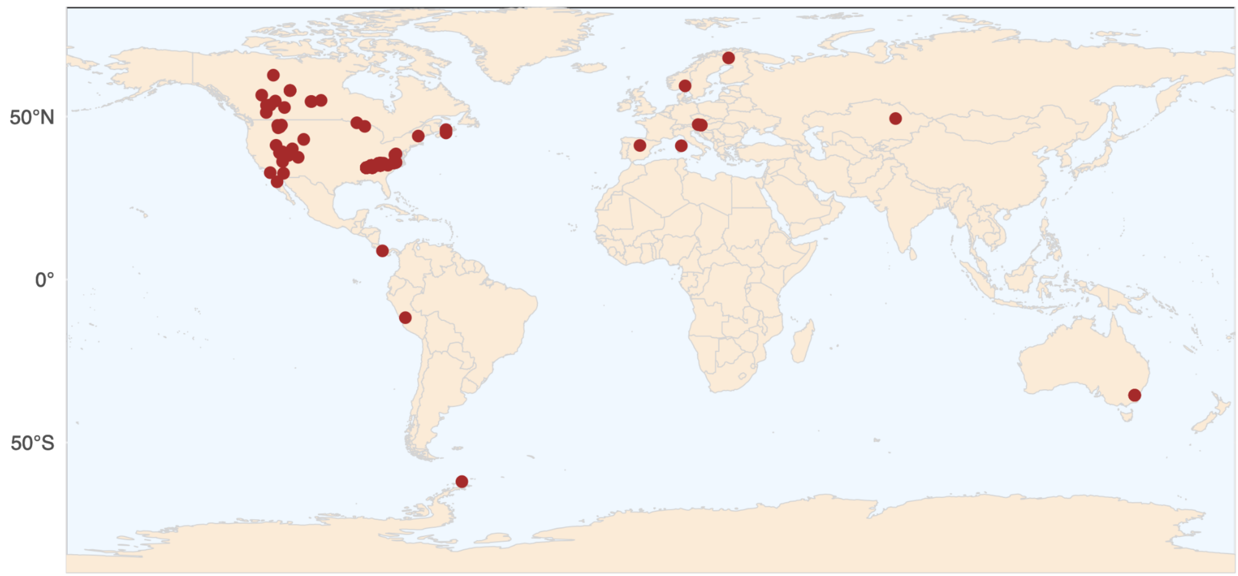


**Extended Data Fig. 2. Sampling locations for the metagenomic data used in the analysis.**

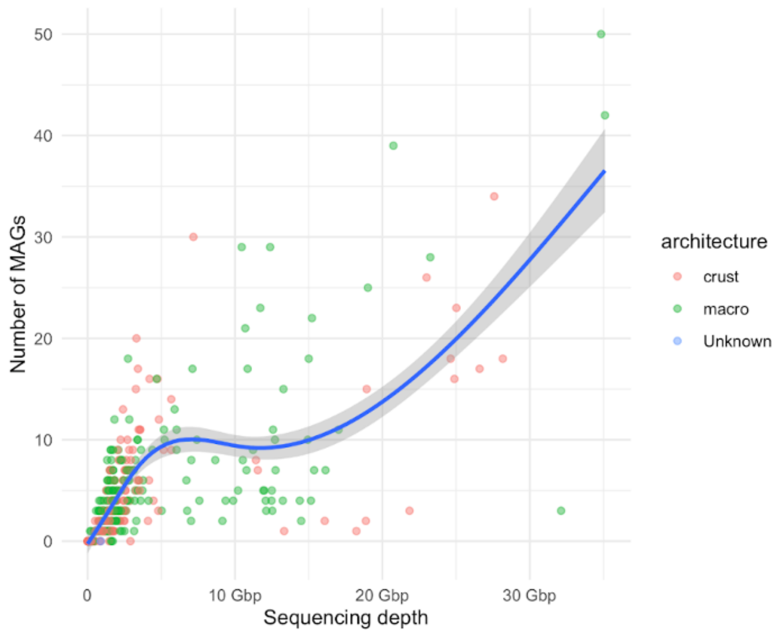


**Extended Data Fig. 3. Number of recovered genomes as a function of sequencing depth (bp).**

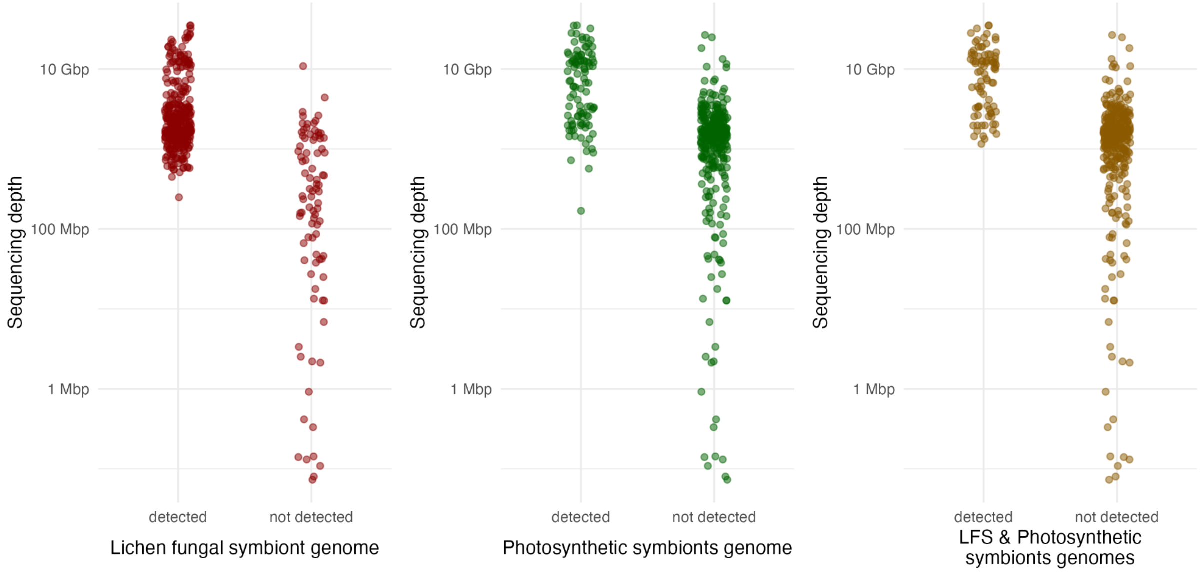


**Extended Data Fig. 4. Recovery of genomes of the main two symbionts as a function of sequencing depth.** These graphs are based on the pre-dereplication set of genomes, each dot represents a metagenome and is positioned based on its sequencing depth and on whether it contained genomes assigned to one or both of the two main partners: A. the LFS (lichen fungal symbiont); B. the photobiont partner; C. both the LFS and the photobiont partner.

Each dot represents a metagenome coloured based on the lichen architecture type. The curve indicates a GAM smoothing.

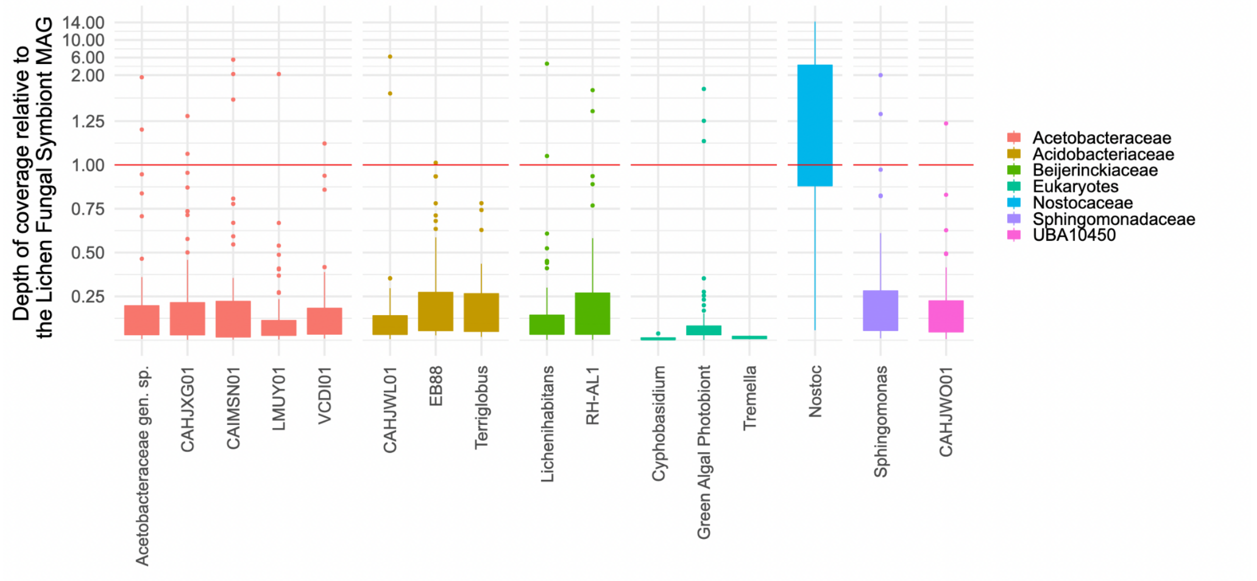


**Extended Data Fig. 5. Relative abundances of symbionts in lichen metagenomes.** The relative abundances were calculated by dividing the coverage depth of the symbiont genome by the coverage of the LFS genome. Here are shown data on the 13 most frequent bacterial genera and the eukaryotes known to be stably associated with lichens. The red line shows 1:1 ratio, where the symbiont is estimated to have the same cellular abundance as the main fungal symbiont.

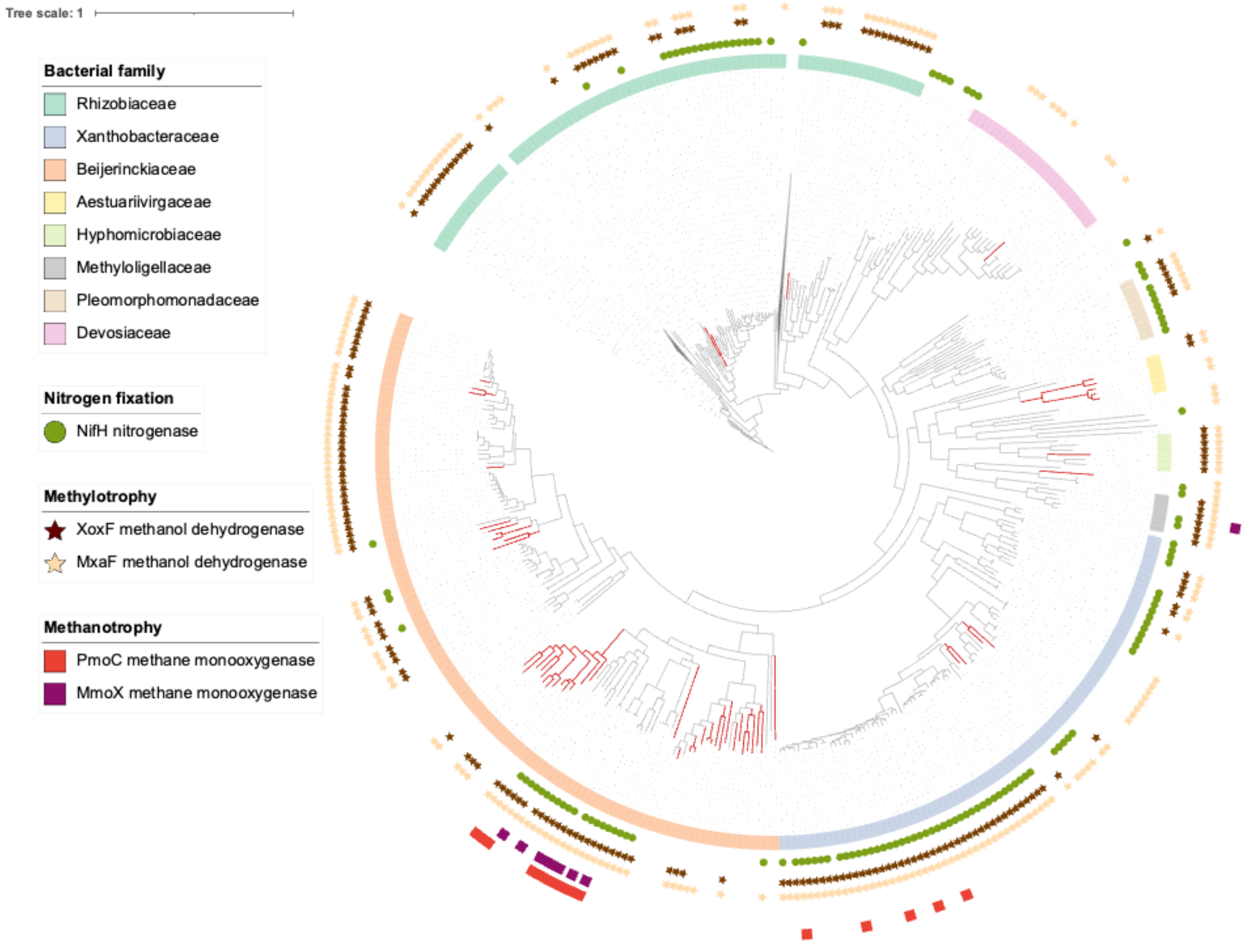


**Extended Data Fig. 6. Maximum likelihood phylogenetic tree of Rhizobiales.** The tree includes published genomes of Rhizobiales and Rhizobiales genomes derived from lichen metagenomes (indicated in red). We generated the alignment of 120 marker genes using GTDB-Tk, and calculated the tree using IQ-TREE. The color represents family-level taxonomic assignment. We used tblastn to search the genomes for key genes involved in nitrogen fixation and C1 metabolism. The presence of these genes is indicated with symbols. The full-size version of the tree in the graphic and Newick formats are available at FigShare (doi.org/10.6084/m9.figshare.21913170.v1).

### Supplementary Figures

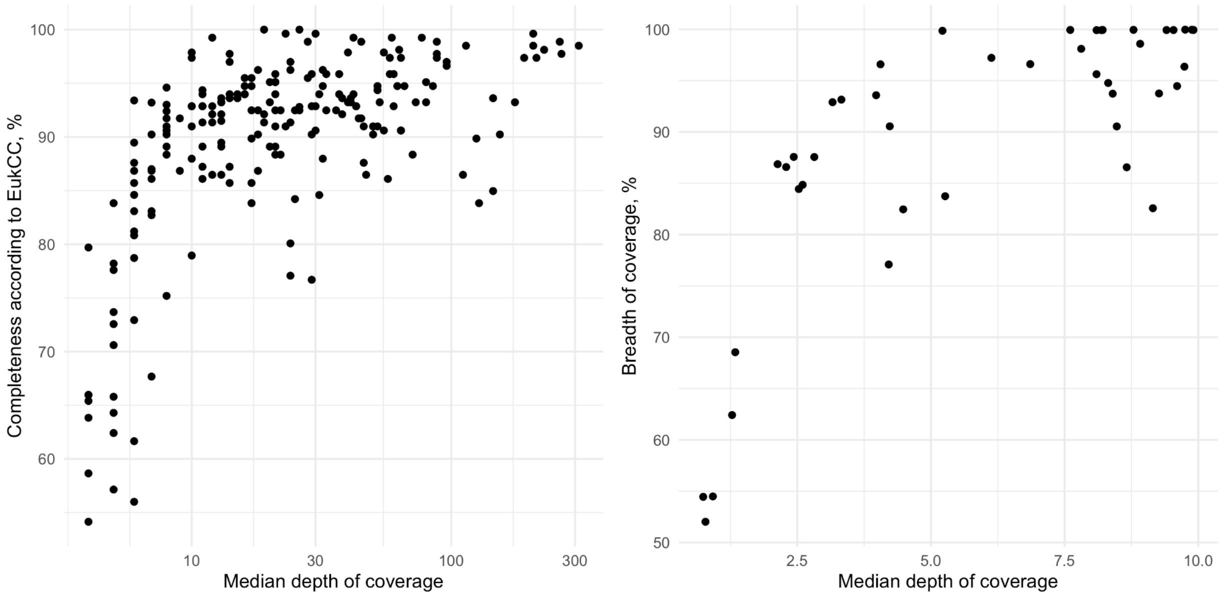


**Supplementary Fig. 1. Genome QC scores as a function of its depth of coverage.** The left panel shows the genome completeness score from EukCC (Saary et al. 2020) as a function of the depth of coverage; the right panel shows the breadth of coverage, as estimated by BWA (Li & Durbin 2009), as a function of the depth of coverage.

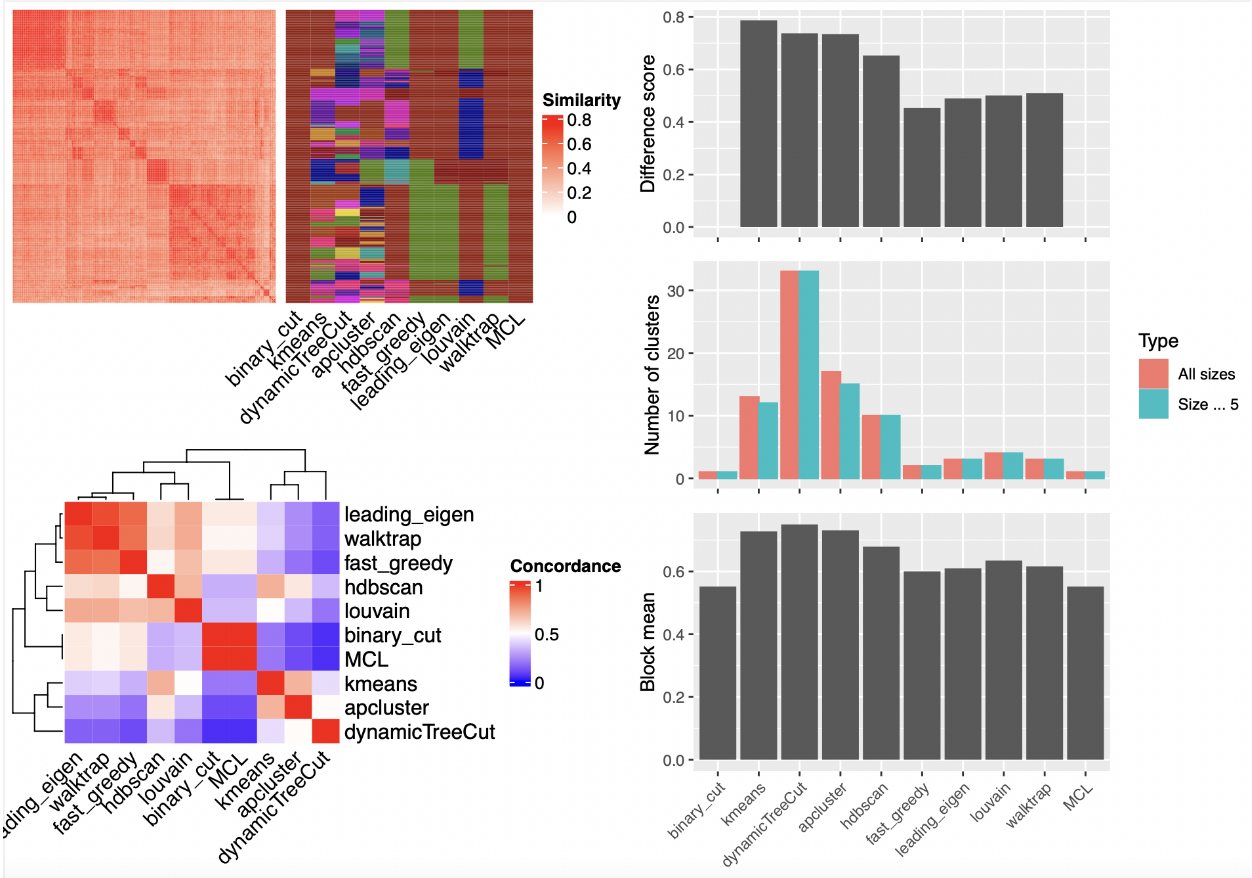


**Supplementary Fig. 2. Comparing different clustering methods for grouping bacterial genomes based on their KEGG profiles.**

We constructed the presence-absence matrix of KEGG modules in the analyzed genomes (all bacterial genomes that met the 90% completeness threshold). Next, we constructed the similarity matrix based on Pearson coefficients and analyzed it using ten clustering methods. The left panel shows: the heatmap of the similarity matrix with different classifications shown as bars (top) and the heatmap of pairwise concordance between the clustering methods (bottom). The right panel shows barplots for each clustering method: Difference score (top, measure of difference between the similarity metric between objects in one cluster and objects in different clusters), Number of clusters (middle), and Block mean (bottom, mean similarity values of the diagonal blocks in the similarity matrix). All the methods were applied to the similarity matrix, which was based on Pearson coefficients. The comparison was done using the SimplifyEnrichment R library.

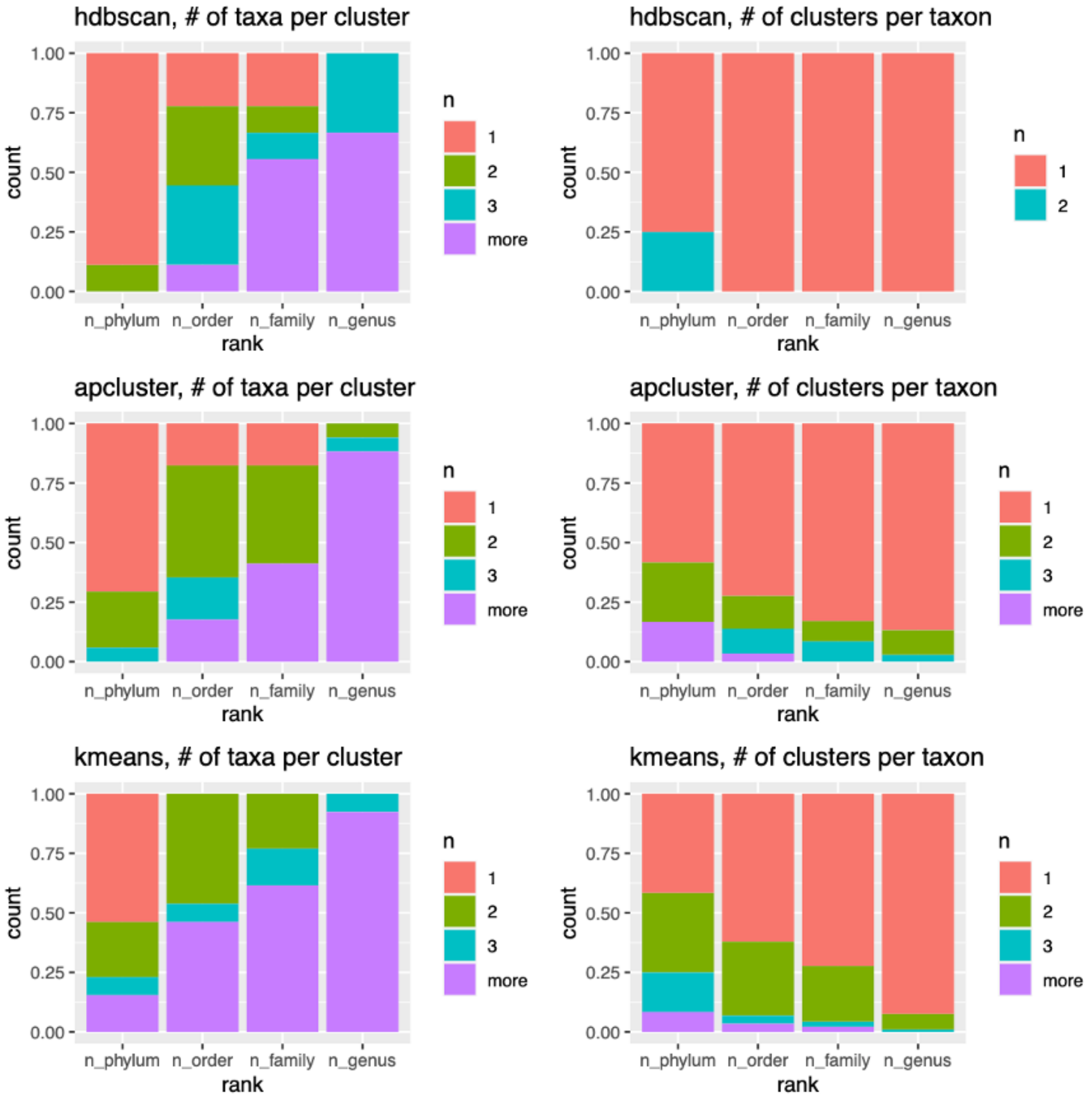


**Supplementary Fig. 3. Taxonomic coherence for the outcomes of three clustering methods: hdbscan, apcluster, and kmeans.**

On the left: percentage of clusters that include 1, 2, 3, or more taxa for each taxonomic rank. On the right: percentage of taxa included in 1, 2, 3, or more different clusters for each taxonomic rank.
